# Structural basis of peptidomimetic agonism revealed by small molecule GLP-1R agonists Boc5 and WB4-24

**DOI:** 10.1101/2022.01.05.475023

**Authors:** Zhaotong Cong, Qingtong Zhou, Yang Li, Li-Nan Chen, Zi-Chen Zhang, Anyi Liang, Qing Liu, Xiaoyan Wu, Antao Dai, Tian Xia, Wei Wu, Yan Zhang, Dehua Yang, Ming-Wei Wang

## Abstract

Glucagon-like peptide-1 receptor (GLP-1R) agonists are effective in treating type 2 diabetes and obesity with proven cardiovascular benefits. However, most of them are peptides and require subcutaneous injection except for orally available semaglutide. Boc5 was identified as the first orthosteric non-peptidic agonist of GLP-1R that mimics a broad spectrum of bioactivities of GLP-1 *in vitro* and *in vivo*. Here, we report the cryo-electron microscopy structures of Boc5 and its analog WB4-24 in complex with the human GLP-1R and G_s_ protein. Bound to the extracellular domain, extracellular loop 2, and transmembrane (TM) helices 1, 2, 3 and 7, one arm of both compounds inserted deeply into the bottom of the orthosteric binding pocket that is usually accessible by peptidic agonists, thereby partially overlapping with the residues A8-D15 in GLP-1. The other three arms, meanwhile, extended to the TM1-TM7, TM1-TM2, and TM2-TM3 clefts showing an interaction feature substantially similar to a previously known small molecule agonist LY3502970. Such a unique binding mode creates a distinct conformation that confers both peptidomimetic agonism and biased signaling induced by non-peptidic modulators at GLP-1R. Further, the conformational difference between Boc5 and WB4-24, two closed related compounds, provides a structural framework for fine tuning of pharmacological efficacy in the development of future small molecule therapeutics targeting GLP-1R.

**Significance:** GLP-1R agonists are efficacious in the treatment of type 2 diabetes and obesity. While most clinically used agents require subcutaneous injection, Boc5, as the first orthosteric non-peptidic agonist of GLP-1R, suffers from poor oral bioavailability that hinders its therapeutic development. The cryo-electron microscopy structures of Boc5 and its closely related analog WB4-24 presented here reveal a previously unknown binding pocket located deeper in the transmembrane domain for non-peptidic GLP-1R agonists. Molecular interaction with this site may facilitate a broad spectrum of in vivo agonistic activities, in addition to that with the upper helical bundles presumably responsible for biased signaling. These findings deepen our understanding of peptidomimetic agonism at GLP-1R and may help design better drug leads against this important target.

## Introduction

It is estimated that diabetes affect the health of more than 10% adults worldwide, with approximately 90% of those suffer from type 2 diabetes mellitus (T2DM) (1). Glucagon-like peptide-1 receptor (GLP-1R) agonists are effective in treating T2DM and obesity, with salient benefits for cardiovascular diseases (2-4). Despite the therapeutic success, this class of medications is suboptimal due to the requirement of subcutaneous injection. With the established therapeutic effects that derive mainly from potentiation of glucose-dependent insulin secretion and suppression of appetite (5), considerable efforts have been made in developing orally available GLP-1R agonists. Semaglutide (Rybelsus®) is the first and only one noninjectable formulation in the clinic in spite of its low bioavailability and frequent gastrointestinal complaints, such as nausea and vomiting (6, 7). To circumvent these problems, search for non-peptidic agonists suitable for oral administration has never stopped.

A number of small molecule GLP-1R agonists in different chemotypes were documented, including quinoxalines, sulfonylthiophenes, pyrimidines, phenylalanine derivatives, substituted cyclobutanes, azoanthracene and oxadiazoanthracene derivatives (8, 9). A few compounds reported in the patent literature have progressed to clinical trials (10). These compounds were reported to mimic GLP-1 to induce insulin release, body weight decrease and cardiovascular disease risk reduction, with variable pharmacokinetic profiles (9). Recently, insulinotropic activity upon oral intake of LY3502970 or PF-06882961 was demonstrated in cynomolgus monkeys (11). Similar result was obtained in mice bearing humanized GLP-1R as well after oral dosing of TTP273 (12).

Boc5, a substituted cyclobutane, was identified as the first orthosteric non-peptidic agonist at GLP-1R (13). Our previous studies show that it is of peptidomimetic nature with dual efficacies in mouse models of T2DM and obesity (14). Pharmacologically, Boc5 was capable of eliciting cAMP accumulation to the same level achieved by GLP-1 and stimulating insulin secretion both *in vitro* and *in vivo* (13). Both chronic oral and intraperitoneal administration of Boc5 in *db/db* mice reduced HbA1c, blood glucose and weight gain (13-15). WB4-24, a closely related analog of Boc5, exhibited more potent *in vitro* and *in vivo* bioactivities (15, 16). While Boc5 and WB4-24 are not “druggable” due to the poor solubility and low oral bioavailability (15), they may represent a unique pharmacophore for optimization. Nonetheless, they could certainly serve as a useful tool to probe the molecular mechanism by which non-peptidic ligands recognize and activate GLP-1R. Here we report the cryo-electron microscopy (cryo-EM) structures of GLP-1R–G_s_ complex bound to Boc5 or WB4-24 to provide valuable insights into the structural basis of their binding mode and signaling properties.

## Results

### Structure Determination

To increase binding of Boc5 and WB4-24 to the GLP-1R–G_s_ complex during purification, the solubility of the compounds was improved by about 1,000 times using a surfactant (poloxamer188, P188) that had no significant impact on receptor activation (*SI Appendix*, Fig. S1A). The complexes were further stabilized by Nb35 and NanoBiT tethering strategy (17, 18), purified using size exclusion chromatography (SEC), and verified by SDS gel (*SI Appendix*, Figs. S1E and S1F). The cryo-EM was used to determine the structures of the complexes bound to Boc5 and WB4-24 at a global nominal resolution of 2.61 Å and 3.09 Å, respectively (Fig. 1 and *SI Appendix*, Figs. S2, S3 and Table S1), with well-defined density for each transmembrane (TM) helix, the compounds, and a majority of the loop regions (*SI Appendix*, Fig. S4). The cryo-EM maps allowed us to build an unambiguous model for most regions of the complexes except for the flexible α-helical domain (AHD) of Gα_s_, the stalk between TM1 and the extracellular domain (ECD) which were poorly resolved in most cryo-EM GPCR–G_s_ complex structures. Extracellular loop (ECL) 3 between V370 and T377, and intracellular loop (ICL) 3 between N388 and D344 were also poorly resolved and not modeled for both structures.

**Fig 1.**
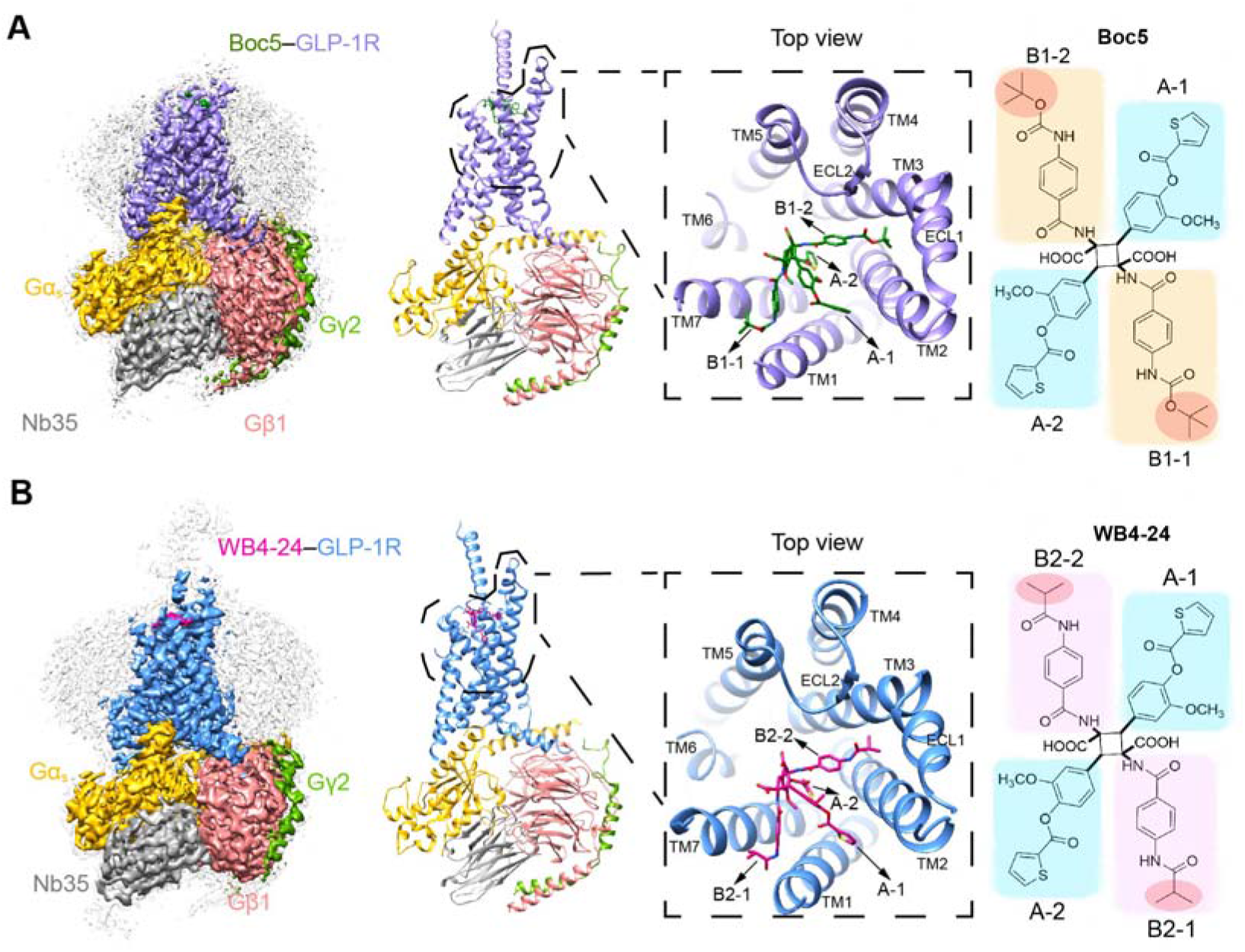
Overall cryo-EM structures of the Boc5- and WB4-24-bound GLP-1R–G_s_ complexes. (*A*) The cryo-EM map with a disc-shaped micelle (left) and cartoon representation (middle) of the Boc5-bound complex. The top view shows the binding pose of Boc5. The right panel shows the chemical structure of Boc5 (molecular weight: 1018) with two 3-methoxy-4-(thiophene-2-carbonyloxy)phenyl groups (blue shade) named A-1 and A-2, respectively, and two 4-((t-butoxycarbonyl)-amino)phenyl groups (orange shade) named B1-1 and B1-2, respectively. (*B*) The cryo-EM map with a disc-shaped micelle (left) and cartoon representation (middle) of the WB4-24-bound complex. The top view shows the binding pose of WB4-24. The right panel shows the chemical structure of WB4-24 (molecular weight: 1074) with two 3-methoxy-4-(thiophene-2-carbonyloxy)phenyl groups (blue shade) that are the same as A-1 and A-2 of Boc5, and two 4-((t-butoxycarbonyl)-amino)phenyl groups (pink shade) named B2-1 and B2-2, respectively. Moieties that are different between Boc5 and WB4-24 are highlighted in red circles. Boc5-bound GLP-1R in purple; WB4-24-bound GLP-1R in dodger blue; Ras-like domain of Gα_s_ in gold; Gβ subunit in deep pink; Gγ subunit in green; Nb35 in gray; Boc5 in dark green; WB4-24 in magenta.

### Receptor Activation

In line with our previous studies (16), Boc5 and WB4-24 displayed a full agonism as GLP-1 at eliciting G_s_-mediated cAMP responses with EC_50_ values of 45 ± 1.12 nM and 23 ± 1.13 nM, respectively (*SI Appendix*, Fig. S1C and Table S2). No detectable β-arrestin recruitment was observed for both Boc5 and WB4-24 (*SI Appendix*, Fig. S1D and Table S2), implying that their glycemic and weight loss effects might be achieved by avoiding β-arrestin mediated receptor desensitization (19).

Compared to the inactive GLP-1R structure (20), WB4-24 binding and G protein coupling facilitated the receptor to undergo significant conformational changes, including the ECD movement toward ECL2, inward movement of the extracellular tip of TM1, outward movement of the extracellular tip of TM2 and the extracellular half of TM7, and subsequent outward movement of the intracellular part of TM6 (Fig. 2A). While similar conformation in the intracellular side was observed in all agonist-bound GLP-1R structures, the extracellular half that agonists directly interact displays ligand-dependent structural features. Compared to the peptidic agonists including GLP-1, the N-terminal α-helix of ECD folded down towards the TMD to stabilize the binding of both WB4-24 and Boc5, whereas the ECL1 reorganized its conformation to accommodate the insertion of the ligands into the TM2-TM3 cleft (Fig. 2A).

**Fig 2.**
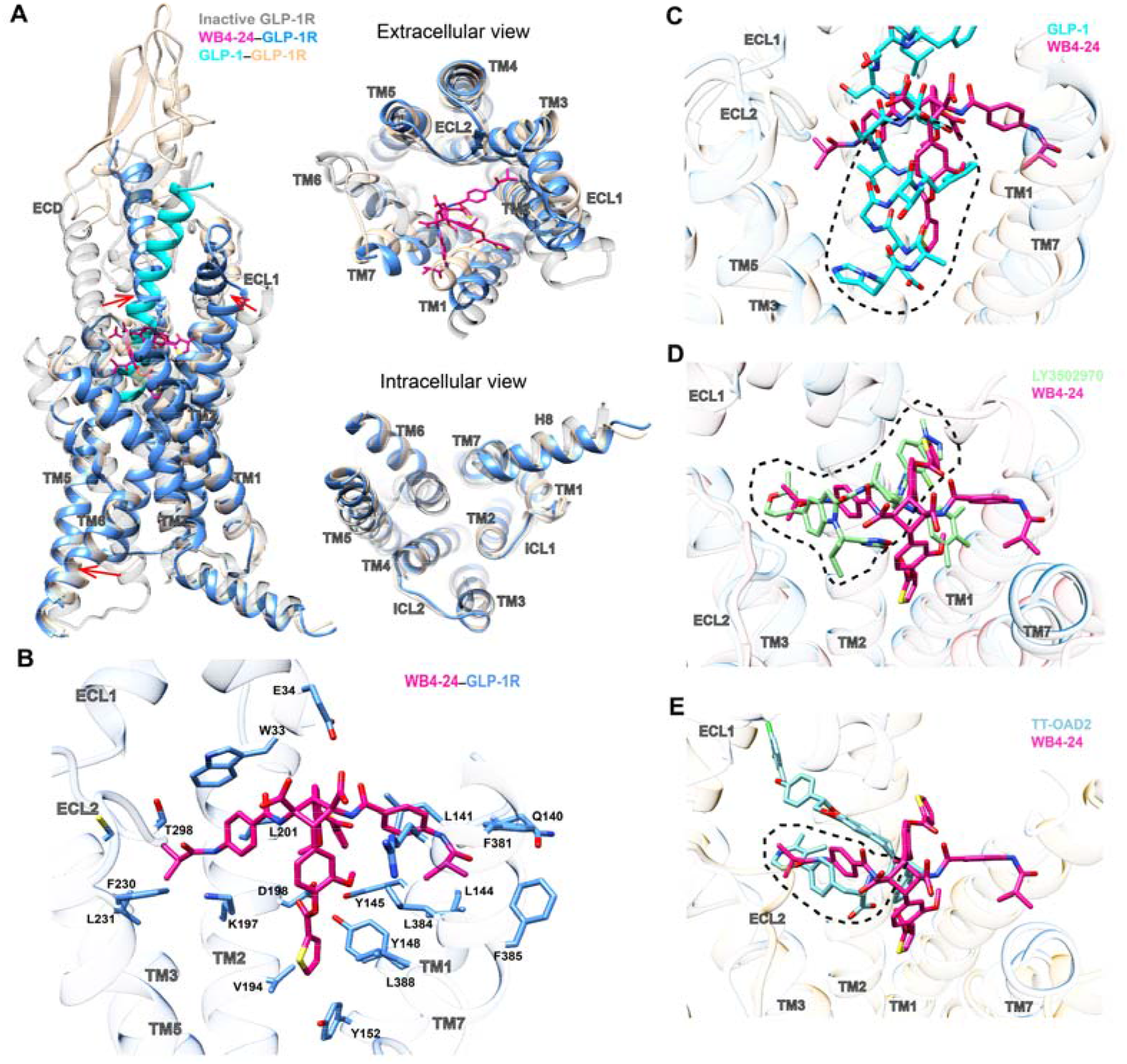
Structural analysis of the WB4-24-bound GLP-1R. (*A*) Structural comparison of the WB4-24-bound GLP-1R with inactive and GLP-1-bound GLP-1R complexes (PDB code: 6×18 and 6LN2). Arrows show the transition from the inactive conformation to WB4-24-induced active state. (*B*) Interactions of WB4-24 within the TM binding cavity. WB4-24 and its interaction residues are shown by sticks. (*C*) Overlay of the WB4-24- and GLP-1-bound receptors reveals that WB4-24 has a partial overlap with the N terminus of GLP-1 in the TM binding cavity. (*D*) Superimposition of the WB4-24- and LY3502970-bound receptors reveals a similar binding pose and a large degree of overlap in the GLP-1R TM cavity. (*E*) Superimposition of the WB4-24- and TT-OAD2-bound receptors reveals a partial overlap in the TM2-TM3 cleft.

WB4-24 anchored in the orthosteric pocket by extensively interacting with residues in TMs 1-3, 7, ECL2 and the N-terminal α-helix through its four arms (labelled in Fig. 1), while the residues in TMs 4-6, ECL1 and ECL3 provided negligible direct contacts (Fig. 2B). Specifically, the 3-methoxy-4-(thiophene-2-carbonyloxy)phenyl group of the compounds (arm A-2, blue shading in Fig. 1) deeply penetrated into the bottom of the orthosteric pocket and partially overlapped with the positions occupied by peptide residues A8^GLP-1^-D15^GLP-1^, but approached to TM1 in a manner closer than that of GLP-1 (Fig. 2C). At the base of the binding pocket of GLP-1-bound GLP-1R, A8^GLP-1^, E9^GLP-1^ and F12^GLP-1^ engaged in an extensive network with TM1 and TM7, including hydrogen bonds with Y152^1.47b^ [class B GPCR numbering in superscript (21)] and R190^2.60b^, hydrophobic interactions with L141^1.36b^, L144^1.39b^ and Y148^1.43b^ in TM1, and E387^7.42b^ and L388^7.43b^ in TM7 (22). A majority of these GLP-1 interacting residues at the binding pocket base also made contact with the thiophene moiety of WB4-24, including L141^1.36b^, L144^1.39b^, Y148^1.43b^, Y152^1.47b^, L384^7.39b^ and L388^7.43b^, despite their distinct chemotypes and binding poses between WB4-24 and GLP-1. Other notable interactions with arm A-2 are the hydrophobic contacts with V194^2.64b^ in TM2, which receives the support of our mutagenesis study where V194A dramatically reduced the efficacy of cAMP responses induced by either Boc5 or WB4-24, with a stronger influence seen with WB4-24-treated cells (Fig. 2B, and *SI Appendix*, Fig. S5 and Table S3). The N-terminal residues H7^GLP-1^ and A8^GLP-1^ that mainly interact with TM3 and TM5 are crucial for receptor activation, as truncation of these two residues, *i*.*e*., GLP-1[9-36]NH_2_, lowers both affinity and efficacy of GLP-1 (23). Limited by the molecular size, both WB4-24 and Boc5 failed to directly interact with the residues in TM3 and TM5 like GLP-1, but they converged to induce a similar conformation of the central polar network at the bottom of the orthosteric pocket. These features may contribute to the peptidomimetic properties of the two ligands.

The other 3-methoxy-4-(thiophene-2-carbonyloxy)phenyl group (arm A-1, blue shading in Fig. 1B) and the 4-((t-butoxycarbonyl)-amino)phenyl group (arm B2-1, pink shading in Fig. 1B) of WB4-24 extended to the TM1-TM2 and TM2-TM3 clefts, respectively. In our molecular dynamics simulation experiments, these two arms steadily interacted with the two clefts (*SI Appendix*, Fig. S6) also occupied by other small molecule agonists, such as LY3502970 (Fig. 2D) and TT-OAD2 (Fig. 2E), where a similar V-shaped orientation within this region was commonly adopted. The 2,2-dimethyl-tetrahydropyran and 4-fluoro-1-methyl-indazole arms of LY3502970 interacted with multiple residues including W33^ECD^, L141^1.36b^, K197^2.67b^, L201^2.71b^, F230^3.33b^ and T298^ECL2^ (Fig. 2D) in a manner similar to WB4-24. However, the 4-((t-butoxycarbonyl)-amino)phenyl group (arm B2-2) of WB4-24 is bulkier than the 3,5-dimethyl-4-fluoro-phenyl ring of LY3502970, thereby conferring distinct interactions with the residues in TM1 and TM7 including Q140^1.35b^, F381^7.36b^ and F385^7.40b^ (Fig. 2B). Together, these specific conformations of the four arms of WB4-24 establish a unique binding mode for this class of molecules.

### Improved Bioactivity

A series of Boc5 analogues were designed to improve bioactivities by varying the length and steric hindrance of R1 groups (red shading in Fig. 1). When the methoxy group of Boc5 (red shading in Fig. 1A) was changed to smaller moieties, the potency of ligand binding and receptor activation were improved, with the tertiary butyl (red shading in Fig. 1B) as the optimum (16). Quantitative analysis of both Boc5- and WB4-24-bound GLP-1R structures may provide mechanistic insights into the increased bioactivity of WB4-24 relative to Boc5.

Structural superimposition reveals that the two molecules adopt a similar binding pose (Fig. 3A), but notable differences in ligand recognition and associated TM bundle conformation were observed (Figs. 3B and 3C). Due to a larger steric hindrance of R1 groups, Boc5 pushed the extracellular tips of TMs 2, 3 and 7 to move outward by 1.6 Å (measured by Cα carbon of Y205^2.75b^), 1.2 Å (measured by Cα carbon of S225^3.28b^) and 2.0 Å (measured by Cα carbon of R380^7.35b^), respectively, compared with WB4-24 (Fig. 3B and 3C). Given the fundamental role of TM6-ECL3-TM7 in GLP-1R-mediated signaling (12, 24), these subtle differences between Boc5 and WB4-24 in TM7 conformation may alter their potencies. Of note, R380^7.35b^ in TM7 formed a salt bridge with the carboxylic group of WB4-24, in a way similar to D15^GLP-1^ in GLP-1 (Fig. 3D). Previous structure-activity relationship (SAR) studies demonstrated that the two carboxylic groups of Boc5 are essential for its bioactivity, whose modification by either amide or ester led to a total loss of activity (16). The importance of R380^7.35b^ was confirmed by alanine mutagenesis that almost abolished the activity of WB4-24 but had a weaker impact on Boc5-induced cAMP production (Fig. 3E and *SI Appendix*, Table S3). Besides the conformational changes at the TM, sidechain orientation of several residues including L144^1.38b^ and K197^2.67b^ were distinct between Boc5 and WB4-24. Alanine mutation of them had a slightly reduced effect on Boc5 than WB4-24 elicited cAMP signaling (Fig. 3E and *SI Appendix*, Table S3). Collectively, these results provide a quantitative measure of pharmacology between two closely related molecules at a near-atomic level.

**Fig 3.**
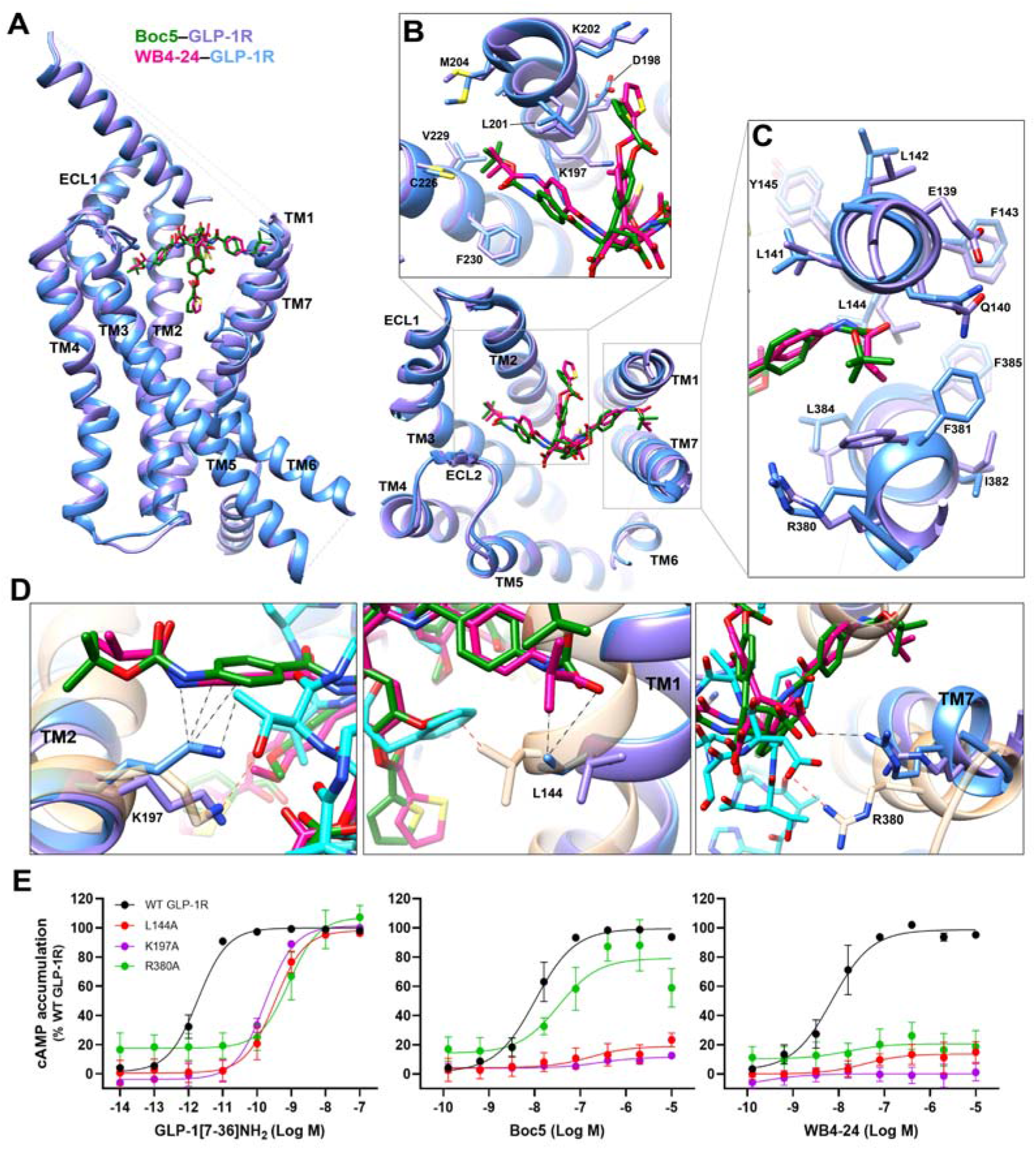
Comparison of GLP-1R conformation and ligand binding pocket stabilized by Boc5 and WB4-24. (*A*) Overlay of Boc5 and WB4-24-bound GLP-1R structures. (*B, C*) Close up of the TM bundles viewed from the extracellular side reveals conformational differences in TM2/TM3 (*B*) and TM1/TM7 (*C*) backbones when stabilized by different arms of Boc5 and WB4-24. (*D*) WB4-24 makes more extensive contacts with TM bundles than Boc5 by interacting with residues L144, K197 and R380 which contribute to the agonism of GLP-1. The interactions of the residues with WB4-24 and GLP-1 are indicated by black and red dashed lines, respectively. (*E*) Mutagenesis analysis of residues L144, K197 and R380 for GLP-1, Boc5 and WB4-24 show the requirement of these contacts for receptor activation. Data are shown as means ± S.E.M from at least three independent experiments.

### Peptidomimetic site

Structural comparison of the GLP-1R structures bound by various small molecule agonists and GLP-1 suggests that Boc5 and WB4-24 mimic the peptidic binding to an extent unseen before. Fig. 4A shows that both compounds penetrated into the GLP-1R TMD pocket deeper than all other non-peptidic ligands reported previously (by 2∼4 Å as measured vertically). Such a binding mode enabled them to directly interact with the bottom of the binding pocket only accessible by peptidic agonists (Fig. 4B-D). Consistently, the interface area between Boc5 and the pocket is 360 Å^2^, almost half of GLP-1 (737 Å^2^), significantly larger than other small molecules (∼250 Å^2^ for LY3502970, CHU-128 and TT-OAD2; 90 Å^2^ for PF-06882961 and 40 Å^2^ for RGT1383). The latter two compounds additionally occupied the cleft between ECL1 and the N-terminal α-helix of ECD, thereby stabilizing the complex by forming multiple strong contacts with ECL1 (Fig. 4B). As a comparison, ECL1 contributed negligible contacts in the cases of Boc5 and WB4-24. Besides this pocket, the TM1-TM7 cleft was also occupied by Boc5 or WB4-24 in a manner more efficient than that of all the other small molecules, supported by the agonist-TM1/7 interface area calculated for Boc5 (680 Å^2^) and WB4-24 (749 Å^2^), respectively, close to that of GLP-1 (890 Å^2^) and remarkably larger than other non-peptidic ligands, especially TT-OAD2 (Fig. 4D). These two newly-identified sites bound by Boc5 and WB4-24 offer valuable clues for the design of next generation peptidomimetic GLP-1R agonist.

**Fig 4.**
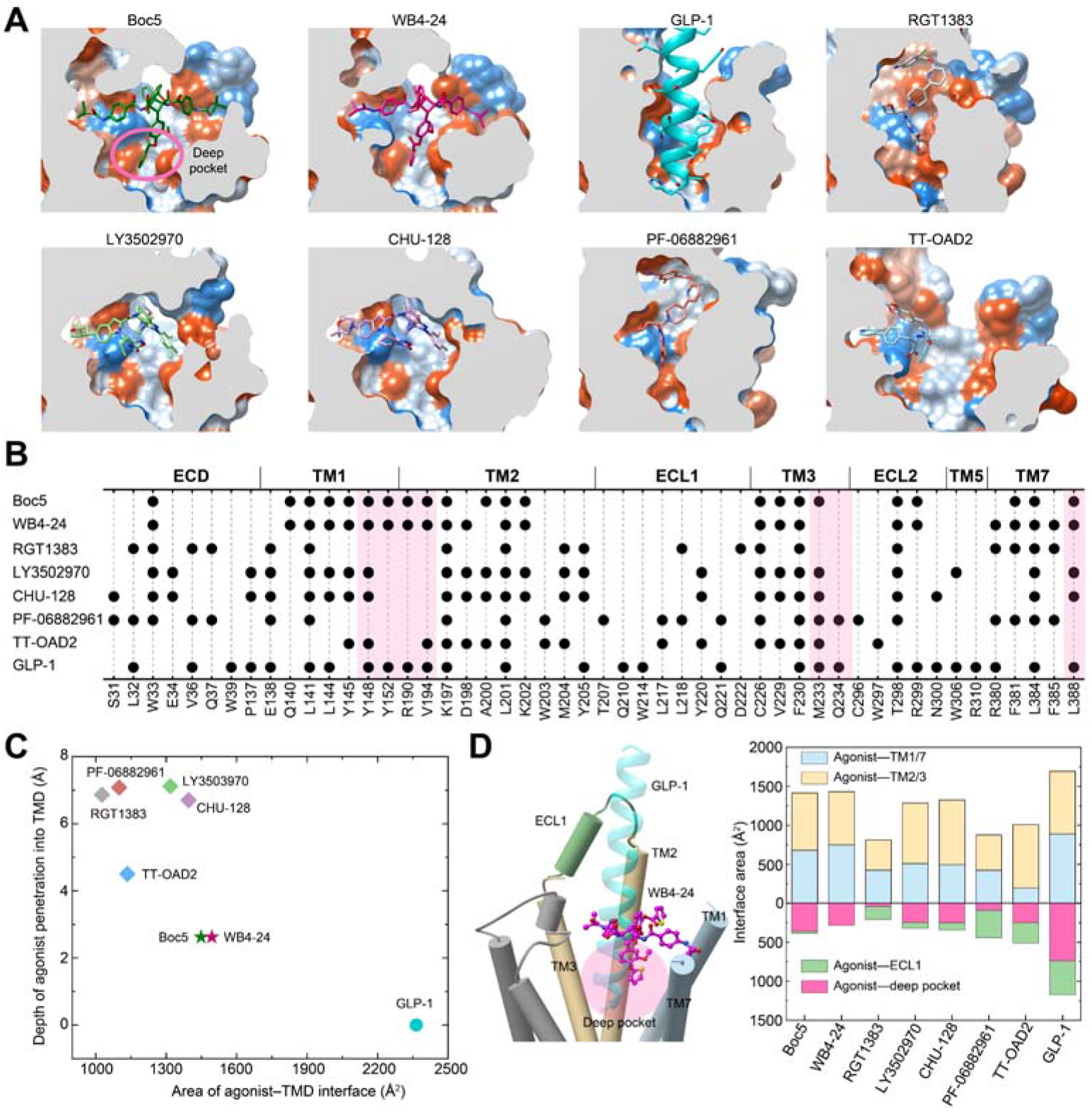
Peptidomimetic binding mode of Boc5 and WB4-24. (*A*) Structural comparison of the binding conformations of peptidic and non-peptidic agonists of GLP-1R. The receptor is shown in surface representation and colored from dodger blue for the most hydrophilic region, to white, and to orange red for the most hydrophobic region. Small molecule agonists and GLP-1 are shown as sticks and ribbon, respectively. (*B*) Schematic diagram of the interactions between peptidic and non-peptidic agonists and GLP-1R. (*C*) Scatter plot of agonist binding in GLP-1R. All structures were superimposed on the GLP-1-bound GLP-1R coordinates from the OPM database (PDB: 6×18) using the TMD Cα atoms. X and Y axis represent the depth of the agonist that penetrated into the GLP-1R TMD pocket and the buried surface area between agonist and TMD, respectively. The former is defined as the minimum vertical distance between the heavy atoms of certain agonist and the OE1 atom of E9^P^ in GLP-1 (the deepest atom of GLP-1 that inserted into the TMD), while the latter was calculated using freeSASA. (*D*) Interface comparison across peptidic and non-peptidic agonists of GLP-1R. Different regions of the GLP-1R TMD pocket including TM1/7, TM2/3, ECL1 (residues S206-D222) and the deep pocket (defined as the collection of the residues Y148^1.43b^-F195^2.65b^, M233^3.36b^-V281^4.57b^, R310^5.40b^-F367^6.56b^ and L388^7.43b^-W420^8.61b^) were subjected to the interface area calculation.

Examination of various molecular recognition patterns of GLP-1R by non-peptidic agonists reveals a common region (the extracellular tips of TM2 and TM3) that constitutes an anchoring platform for agonist binding. Specifically, the sidechains of K197^2.67b^ and F230^3.33b^ pointed to the TMD core and uplifted a small molecule, allowing the latter to be further covered by the folded-down N-terminal α-helix of ECD, particularly W33^ECD^ (Fig. 4B). The TM1-TM2 cleft to which the A-1 branch of Boc5 or WB4-24 inserted was also touched by CHU-128, LY3502970 and TT-OAD2 but neither RGT1383 nor PF-06882961. It was reported that lipidated K10 in peptide 20 and positive allosteric modulator LSN3160440 bound exactly to this cleft via formation of more interactions with receptor (25), indicating a potential region for lead optimization. Similar phenomena were also observed in the TM2-TM3 cleft, to which all other non-peptidic ligands (TT-OAD2 in particular) except RGT1383 and PF-06882961 penetrated using one of the arms. Despite a deficiency in making contacts with the above two regions (TM1-TM2 and TM2-TM3 clefts), RGT1383 and PF-06882961 successfully triggered an inward movement of ECL3 and the extracellular end of TM7, consequently creating a tightly packed orthosteric pocket to accommodate agonist binding (Fig. 4A, B). These analyses highlight the diversity in signaling initiation by non-peptidic ligands.

## Discussion

Boc5 was identified as the first non-peptidic GLP-1R agonist with a broad spectrum of *in vitro* and *in vivo* bioactivities as GLP-1 (13-15). To explore the SAR of this compound, over fifty derivatives with various modifications on the cyclobutane core and four arms (16) were synthesized and evaluated. Among them, WB4-24 stood out as the most potent analog of Boc5 (15, 16, 26). However, suffering from low oral bioavailability due to metabolism by esterase in the gut as well as limitation of photosynthesis (15), this class of GLP-1R agonists is precluded from further development. Meanwhile, a handful of non-peptidic ligands with distinct pharmacological properties were identified in the past decades including TT-OAD2 (12), LY3502970 (27), CHU-128 (22), PF-06882961 (22) and RGT1383 (28). While a few have entered into clinical trials, none was regulatory approved for clinical use.

The cryo-EM structures of GLP-1R in complex with Boc5 or WB4-24 presented here expanded our knowledge of ligand recognition and receptor activation from three aspects. First, the unique binding mode of Boc5 and WB4-24 whose one arm (A-2) inserted deeply into the bottom of the orthosteric pocket usually accessible only by peptidic agonists reveals a previously unknown interaction site for small molecule agonists. Second, quantitative analysis between Boc5- and WB4-24-bound GLP-1R structures, together with the mutagenesis data, provide mechanistic insights into of the improved bioactivity of WB4-24 over Boc5 at the near-atomic level. Third, our systematic review of all the reported GLP-1R structures bound by non-peptidic ligands (Fig. 5) uncovers both common and unique features of ligand recognition distinctively related to the pharmacological property of each other. The extracellular tips of TM2 and TM3 are one common interacting region that make extensive contacts with all bound ligands regardless of peptidic or non-peptidic nature. Meanwhile, individual ligands may also have their specific interaction sites: peptides including GLP-1 have rich contacts with the bottom of the orthosteric pocket (the deep pocket) and the ECL1 but not the TM1-TM2 or TM2-TM3 cleft; TT-OAD2 steadily inserts into the TM2-TM3 cleft while CHU-128 and LY3502970 additionally occupy the TM1-TM2 cleft; both PF-06882961 and RGT1383 alternatively chose the ECL1 and fold the ECL3 and the extracellular tip of TM7 inward as interaction sites. Surprisingly, Boc5 and WB4-24 having four long-extended arms simultaneously occupy the deep pocket, TM1-TM2, TM2-TM3 and TM1-TM7 clefts, allowing them to achieve peptidomimetic agonism and consequently show GLP-1-like *in vivo* activities such as insulin release, appetite suppression, HbA1c reduction and weight loss, *etc*. (*SI Appendix*, Table S5).

**Fig 5.**
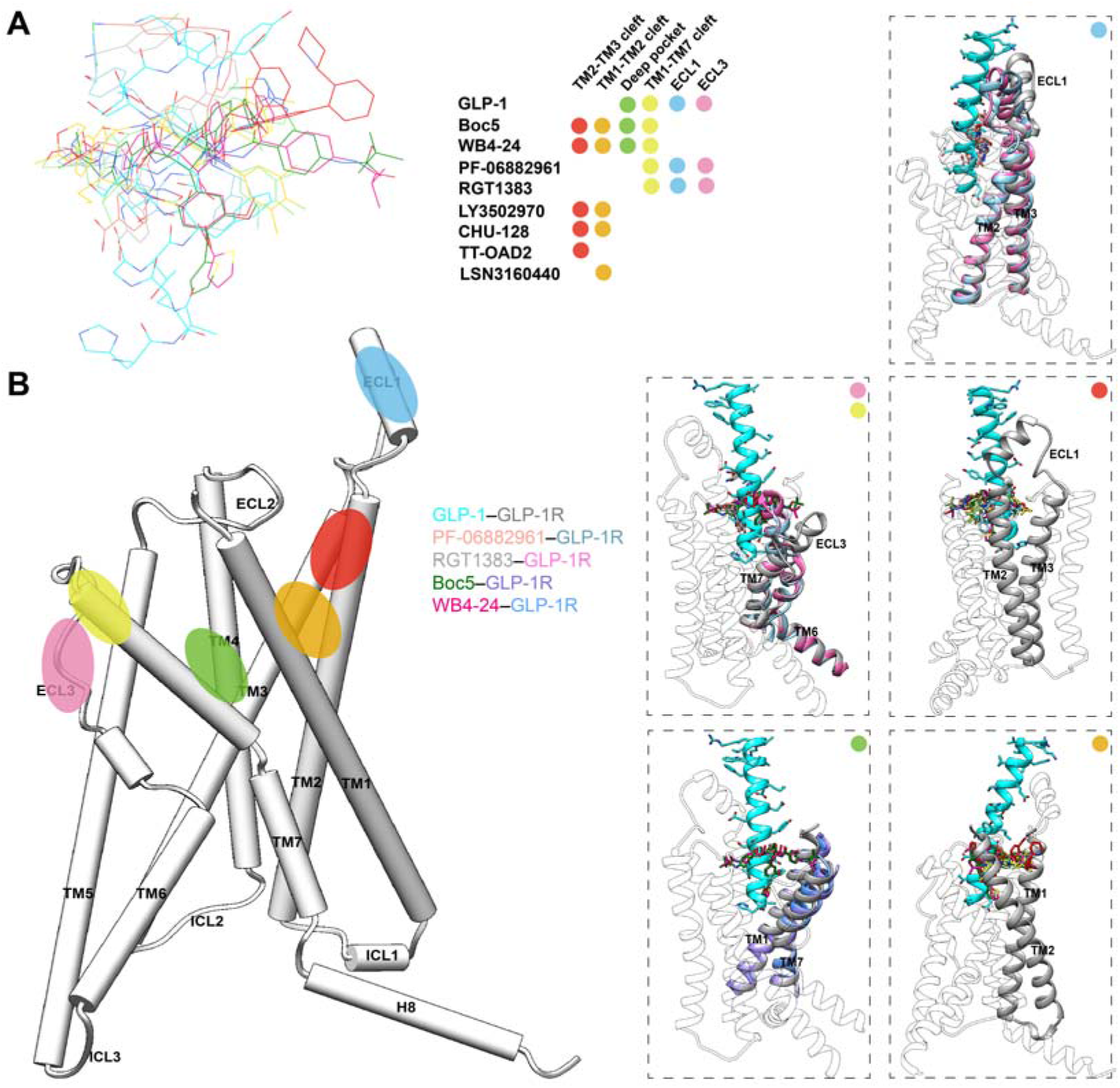
Systematic comparison of ligand recognition patterns of different non-peptidic GLP-1R agonists. (*A*) Binding poses of the reported non-peptidic agonists, GLP-1 (residues H7-L20 is shown) and a positive allosteric modulator LSN3160440. The binding sites of these ligands mainly locate in five regions represented by color dots, including the clefts of TM1-TM2 (orange), TM2-TM3 (red) and TM1-TM7 (yellow), ECLs 1 (blue) and 3 (pink), and the deep orthosteric pocket (green). (*B*) Positions of the binding sites identified among published GLP-1R structures. The dashed boxes show the binding modes of the agonists in specific regions. Color is consistent with binding sites in panel (*A*).

These analyses provide a molecular basis of pharmacological consequences rendered by GLP-1R agonists of different chemical nature. It is known that TM7 conformation is important for peptidomimetic properties of small molecule agonists (27). In the PF-06882961-bound structure, TM7 moved inward dramatically, forming a similar polar interaction network observed in the GLP-1-bound structure. Thus, PF-06882961 was able to better mimic the pharmacological effects of GLP-1 than their counterparts reported in the literature (*SI Appendix*, Table. S5). Such an inward movement is followed by TM6 to form an extensive polar interaction network involving R380^7.35^, thereby constituting the conformation of TM6-ECL3-TM7. In addition, contraction of TM7 induced by PF-06882961 moves F385^7.40^ to the TM1-TM7 cleft to stabilize overall sidechain rearrangements among key residues located in TMs 1 and 2 and mediated by residues Y148^1.43^, K197^2.67^ and D198^2.68^ (22). Intriguingly, in the Boc5- and WB4-24-bound structures, the interactions with TM1, TM2 and TM7 are mainly made by A-1 and A-2 arms, supporting an essential role played by thiophene groups in receptor activation. Removal of them in A-2 would impair the interaction with TM7, resulting an inactive metabolite M-2 (15). Moreover, compared to full agonists, ligands that exhibit biased agonism towards G protein-dependent cAMP signaling are associated with a distinct conformation of TM6-ECL3-TM7, such as LY35029070 (27), TT-OAD2 (12) and CHU-128 (22). Although the interaction with TM7 in the WB4-24-bound GLP-1R was similar to that in GLP-1-or PF-06882961-bound structures, its contact with ECL3 was not observed, which may account for the biased agonism of Boc5 and WB4-24. Overall, these results imply that stabilizing the conformation of the TM6-ECL3-TM7 region would be required for unbiased as well as peptidomimetic agonism at GLP-1R. Clearly, the newly-identified binding pocket of Boc5 and WB4-24 as well as their interaction mode with the receptor provide valuable clues for developing next generation of non-peptidic modulators with better therapeutic profiles.

## Methods

### Cell culture

*Spodoptera frugiperda* (*Sf*9) insect cells (Expression Systems) were grown in ESF 921 serum-free medium (Expression Systems) at 27°C and 120 rpm. HEK293T cells (American Type Culture Collection) were cultured in Dulbecco’s modified Eagle’s medium (DMEM, Life Technologies) supplemented with 10% fetal bovine serum (FBS, Gibco) and maintained in a humidified chamber with 5% CO_2_ at 37°C.

### Construct preparation

The human GLP-1R was modified with its native signal sequence (M1-P23) replaced by the haemagglutinin (HA) signal peptide to facilitate receptor expression. To obtain a GLP-1R–G_s_ complex with good homogeneity and stability, we used the NanoBiT tethering strategy, in which the C terminus of rat Gβ1 was linked to HiBiT subunit with a 15-amino acid polypeptide (GSSGGGGSGGGGSSG) linker and the C terminus of GLP-1R was directly attached to LgBiT subunit followed by a TEV protease cleavage site and a double MBP tag. An engineered G_s_ construct (G112) was designed based on mini-G_s_ as previously described (29, 30) and was used to purify the Boc5 or WB4-24 bound GLP-1R–G_s_ complex. The constructs were cloned into both pcDNA3.1 and pFastBac vectors for functional assays in mammalian cells and protein expression in insect cells, respectively. All modifications of the receptor had no effect on ligand binding and receptor activation (*SI Appendix*, Fig. S1B and Table S3). Other constructs including the wild-type (WT) and GLP-1R mutants were cloned into pcDNA3.1 vector for cAMP accumulation and whole cell binding assays.

### Solubility analysis

The excess amount of Boc5 and WB4-24 were dissolved by shaking at 37°C and 180 rpm for 72 h in normal saline, 5 mM HEPES, 20% PEG400 or 2% P188, respectively. After centrifugation at 13,000 rpm for 10 min, the supernatant was extracted and diluted to an appropriate concentration with methanol. The saturated solubility of the compounds was determined using high-performance liquid chromatography (HPLC) and estimated by fitting the data to the standard concentration curves.

### Formation and purification of complex

The Bac-to-Bac Baculovirus Expression System (Invitrogen) was used to generate high-titer recombinant baculovirus for GLP-1R-LgBiT-2MBP, G112, Gβ1-HiBiT and Gγ2. P0 viral stock was produced by transfecting 5 μg recombinant bacmids into *Sf*9 cells (2.5 mL, density of 1 million cells per mL) for 96 h incubation and then used to produce P1 and P2 baculoviruses. GLP-1R-LgBiT-2MBP, DNGαs, Gβ1-HiBiT and Gγ2 were co-expressed at multiplicity of infection (MOI) ratio of 1:1:1:1 by infecting *Sf*9 cells at a density of 3 million cells per mL with P2 baculovirus (viral titers>90%). Culture was harvested by centrifugation for 48 h post infection and cell pellets were stored at -80°C until use.

To prepare Boc5 or WB4-24 solution, the compound was first dissolved in DMSO with a final concentration of 100 mM, and then distributed by 2% P188 solution (1:10, v/v) under ultrasonic condition at room temperature (RT) until a clear solution was obtained. The cell pellets were thawed and lysed in a buffer containing 20 mM HEPES, pH 7.5, 100 mM NaCl, 10% (v/v) glycerol, 10 mM MgCl_2_, 1 mM MnCl_2_ and 100 μM TCEP supplemented with EDTA-free protease inhibitor cocktail (Bimake) by dounce homogenization. The complex formation was initiated by the addition of Boc5 or WB4-24 solution (1:100, v/v), 10 μg/mL Nb35 and 25 mU/mL apyrase (New England Bio-Labs). After 1.5 h incubation at RT, the membrane was solubilized in the buffer above supplemented with 0.5% (w/v) lauryl maltose neopentyl glycol (LMNG, Anatrace) and 0.1% (w/v) cholesterol hemisuccinate (CHS, Anatrace) for 2 h at 4°C. The supernatant was isolated by centrifugation at 65,000 *g* for 30 min and incubated with amylose resin (New England Bio-Labs) for 2 h at 4°C. The resin was then collected by centrifugation at 500 *g* for 10 min and washed in gravity flow column (Sangon Biotech) with five column volumes of buffer containing 20 mM HEPES (pH 7.5), 100 mM NaCl, 10% (v/v) glycerol, 5 mM MgCl_2_, 1 mM MnCl_2_, 25 μM TCEP, and 0.1% (w/v) LMNG-0.02% (w/v) CHS supplemented with 10 μM Boc5 or WB4-24, followed by washing with fifteen column volumes of buffer containing 20 mM HEPES (pH 7.5), 100 mM NaCl, 10% (v/v) glycerol, 5 mM MgCl_2_, 1 mM MnCl_2_, 25 μM TCEP, and 0.03% (w/v) LMNG-0.01% (w/v) glyco-diosgenin (GDN, Anatrace)-0.008% (w/v) CHS supplemented with 10 μM Boc5 or WB4-24. The protein was then incubated overnight with TEV protease (customer-made) on the column to remove the C-terminal 2MBP-tag in the buffer above at 4°C. The flow through was collected next day and concentrated with a 100 kDa molecular weight cut-off concentrator (Millipore). The concentrated product was loaded onto a Superdex 200 increase 10/300 GL column (GE Healthcare) with running buffer containing 20 mM HEPES (pH 7.5), 100 mM NaCl, 10 mM MgCl_2_, 100 μM TCEP, and 0.00075% LMNG-0.00025% GDN-0.0002% (w/v) CHS supplemented with 2 μM Boc5 or WB4-24. The fractions for monomeric complex were collected and concentrated to 15-20 mg/mL for cryo-EM examination.

### Expression and purification of Nb35

Nb35 with a C-terminal 6 × His-tag was expressed in the periplasm of *E. coli* BL21 (DE3), extracted and purified by nickel affinity chromatography as previously described (31). The HiLoad 16/600 Superdex 75 column (GE Healthcare) was used to separate the monomeric fractions of Nb35 with running buffer containing 20 mM HEPES, pH 7.5 and 100 mM NaCl. The purified Nb35 was flash frozen in 30% (v/v) glycerol by liquid nitrogen and stored in -80°C until use.

### Cryo-EM data acquisition

The concentrated sample (3.5 μL) was applied to glow-discharged holey carbon grids (Quantifoil R1.2/1.3, 300 mesh), and subsequently vitrified using a Vitrobot Mark IV (ThermoFisher Scientific) set at 100% humidity and 4°C. Cryo-EM images were collected on a Titan Krios microscope (FEI) equipped with Gatan energy filter and K3 direct electron detector. The microscope was operated at 300 kV accelerating voltage, at a nominal magnification of 46,685× in counting mode, corresponding to a pixel size of 1.071 Å. The total exposure time was set to 7.2 s with intermediate frames recorded every 0.2 s, resulting in an accumulated dose of 80 electrons per Å^2^ with a defocus range of -1.2 to -2.2 μm. Totally, 10,654 images for Boc5–GLP-1R–G_s_ and 10,148 images for WB4-24−GLP-1R−G_s_ complexes were collected.

### Image processing

Dose-fractionated image stacks were subjected to beam-induced motion correction using MotionCor2.1(32). A sum of all frames, filtered according to the exposure dose, in each image stack was used for further processing. Contrast transfer function (CTF) parameters for each micrograph were determined by Gctf v1.06 (33). Particle selection, 2D classifications, ab initio model generation and homogeneous refinement were performed on cryoSPARC.

For the Boc5–GLP-1R–G_s_ complex, template-picking yielded 6,785,990 particles that were subjected to 3D classifications with mask on the receptor to discard false positive particles or particles categorized in poorly defined classes, producing particle projections for further processing. This subset of particles was subjected to further 3D auto-refinement with mask on the complex. A selected subset containing 610,843 projections was subsequently subjected to 3D refinement and Bayesian polishing with a pixel size of 1.045. After last round of refinement, the final map has an indicated global resolution of 2.61 Å at a Fourier shell correlation (FSC) of 0.143. Local resolution was determined using the Bsoft package with half maps as input maps (34).

For the WB4-24−GLP-1R−G_s_ complex, a total of 8,405,468 particles were picked using template-picker. The 3D classifications were performed with mask on the receptor to discard false positive particles or particles categorized in poorly defined classes, producing 3,866,515 particle projections for further processing. This subset of particles was subjected to further 3D auto-refinement with mask on the complex, which were subsequently subjected to a round of 3D classifications with mask on the ECD. A data set of 747,282 particles was subjected to 3D refinement, yielding a final map with a global nominal resolution at 3.09 Å by the 0.143 criteria of the gold-standard FSC. Half-reconstructions were used to determine the local resolution of each map.

### Model building and refinement

The structures of the CHU-128–GLP-1R–G_s_ (22) (PDB: 6×19) was used as an initial template for model building of the Boc5 or WB4-24-bound complexes. Ligand coordinates and geometry restraints were generated using phenix.elbow. Models were docked into the EM density map using UCSF Chimera. This starting model was then subjected to iterative rounds of manual adjustment and automated refinement in Coot (35) and Phenix (36), respectively. The final refinement statistics were validated using the module comprehensive validation (cryo-EM) in PHENIX. Structural figures were prepared in Chimera, Chimera X and PyMOL (https://pymol.org/2/). The final refinement statistics are provided in *SI Appendix*, Table S1.

### cAMP accumulation assay

Peptide-or small molecule-stimulated cAMP accumulation was measured by a LANCE Ultra cAMP kit (PerkinElmer). Briefly, after 24 h transfection with various constructs, HEK293T cells were digested by 0.2% (w/v) EDTA and washed once with Dulbecco’s phosphate buffered saline (DPBS). Cells were then resuspended with stimulation buffer (Hanks’ balanced salt solution (HBSS) supplemented with 5 mM HEPES, 0.5 mM IBMX and 0.1% (w/v) BSA, pH 7.4) to a density of 0.6 million cells per mL and added to 384-well white plates (3,000 cells per well). Different concentrations of ligand in stimulation buffer were added and the stimulation lasted for 40 min at RT. The reaction was stopped by adding 5 μL Eu-cAMP tracer and ULight-anti-cAMP. After 1 h incubation at RT, the plate was read by an Envision plate reader (PerkinElmer) to measure TR-FRET signals (excitation: 320 nm, emission: 615 nm and 665 nm). A cAMP standard curve was used to convert the fluorescence resonance energy transfer ratios (665/615 nm) to cAMP concentrations.

### Whole cell binding assay

HEK293T cells were seeded into 96-well plates (PerkinElmer) coated with poly-D-lysine hydrobromide (Sigma-Aldrich) at a density of 30,000 cells per well and incubated overnight. After 24 h transfection, cells were washed twice and incubated with blocking buffer (DMEM supplemented with 33 mM HEPES, and 0.1% (w/v) BSA, pH 7.4) for 2 h at 37°C. Then, radiolabeled ^125^I-GLP-1 (30 pM, PerkinElmer) and increasing concentrations of unlabeled ligand were added and competitively reacted with the cells in binding buffer (PBS supplemented with 10% (w/v) BSA, pH 7.4) at RT for 3 h. After that, cells were washed with ice-cold PBS and lysed by 50 μL lysis buffer (PBS supplemented with 20 mM Tris-HCl and 1% (v/v) Triton X-100, pH 7.4). Finally, 150 μL of scintillation cocktail (OptiPhase SuperMix, PerkinElmer) was employed and radioactivity (counts per minute, CPM) determined by a scintillation counter (MicroBeta2 plate counter, PerkinElmer).

### β**-arrestin 1/2 recruitment assay**

HEK293T cells were seeded at a density of 30,000 cells per well into 96-well culture plates pretreated with poly-D-lysine hydrobromide. After incubation for 24 h to reach 80% confluence, the cells were transiently transfected with HA-GLP-1R-Rluc8 and β-arrestin 1/2-Venus at a 1:9 mass ratio using lipofectamine 3000 reagent (Invitrogen) and cultured for another 24 h. Thereafter, cells were washed once and incubated for 30 min at 37°C with HBSS buffer (pH 7.4) supplemented with 0.1% BSA and 10 mM HEPES. Five micromolars of coelenterazine h (Yeasen Biotech) was then added and incubated for 5 min in the dark. The bioluminescence resonance energy transfer (BRET) signals were detected with an Envision plate reader by calculating the ratio of emission at 535 nm over emission at 470 nm. A 1.5-min baseline of BRET measurement was taken before the addition of ligand and BRET signal was measured at 10-s intervals for further 9 min. After removing baseline and background readings by subtracting average values of the baseline measurement and average values of vehicle-treated samples, respectively, the AUC across the time-course response curve was determined. Concentration-response curves were plotted using the total area-under-the-curve during the time of measurement post ligand addition.

### Molecular dynamics simulation

Molecular dynamics simulation studies were performed using Gromacs 2020.1. The Boc5–GLP-1R and WB4-24–GLP-1R complexes were built based on the cryo-EM structures and prepared by Protein Preparation Wizard (Schrodinger 2017-4) with G protein and Nb35 nanobody removed. The missing ECD and loops in the GLP-1R were generated by molecular superposition, using USCF Chimera, of the corresponding regions in the CHU-128–GLP-1R–G_s_ complex (22) (PDB: 6×19). The receptor was capped with acetyl and methylamide. All titratable residues were left in their dominant state at pH 7.0. The ligand bound GLP-1R complexes were embedded in a bilayer composed of 211 POPC lipids and solvated with 0.15 M NaCl in explicit TIP3P waters using CHARMM-GUI Membrane Builder v3.5 (37). The CHARMM36-CAMP force filed (38) was adopted for protein, lipids and salt ions. Boc5 and WB4-24 were modeled with the CHARMM CGenFF small-molecule force field (39), program version 2.5. The Particle Mesh Ewald (PME) method was used to treat all electrostatic interactions beyond a cut-off of 12 Å and the bonds involving hydrogen atoms were constrained using LINCS algorithm (40). The complex system was first relaxed using the steepest descent energy minimization, followed by slow heating of the system to 310 K with restraints. Finally, 1,000 ns production simulations without restraints were carried out, with a time step of 2 fs in the NPT ensemble at 310 K and 1 bar using the v-rescale thermostat and the semi-isotropic Parrinello-Rahman barostat (41), respectively.

### Statistical analysis

All functional study data were analyzed using Prism 7 (GraphPad) and presented as means ± S.E.M. from at least three independent experiments. Concentration-response curves were evaluated with a three-parameter logistic equation. The significance was determined with either two-tailed Student’s *t*-test or one-way ANOVA, and P<0.05 was considered statistically significant.

### Data availability

The atomic coordinates and the electron microscopy maps have been deposited in the Protein Data Bank (PDB) under accession codes: xxx (Boc5–GLP-1R–G_s_ complex) and xxx (WB4-24–GLP-1R–G_s_ complex), and Electron Microscopy Data Bank (EMDB) accession codes: xxx (Boc5–GLP-1R–G_s_ complex), and xxx (WB4-24–GLP-1R–G_s_ complex), respectively. All relevant data are available from the authors and/or included in the manuscript or Supplementary Information.

## Acknowledgments

We are indebted to Eric H. Xu, Honglei Ma, Peng-Fei Lan and Ming Lei for valuable discussions. This work was partially supported by National Natural Science Foundation of China 81872915 (M.-W.W.), 82073904 (M.-W.W.), 82121005 (D.Y.), 81973373 (D.Y.) and 21704064 (Q.Z.); National Science & Technology Major Project of China–Key New Drug Creation and Manufacturing Program 2018ZX09735–001 (M.-W.W.) and 2018ZX09711002–002–005 (D.Y.); National Science and Technology Major Project of China – Innovation 2030 for Brain Science and Brain-Inspired Technology 2021ZD0203400 (Q.Z.); the National Key Basic Research Program of China 2018YFA0507000 (M.-W.W.); Novo Nordisk-CAS Research Fund grant NNCAS-2017–1-CC (D.Y.) and SA-SIBS Scholarship Program (D.Y.). The cryo-EM data were collected at Cryo-Electron Microscopy Research Center, Shanghai Institute of Materia Medica.

## References

1. Reed J, Bain S, & Kanamarlapudi V (2021) A review of current trends with type 2 diabetes epidemiology, aetiology, pathogenesis, treatments and future perspectives. Diabetes Metab Syndr Obes 14:3567–3602.

2. Saraiva FK & Sposito AC (2014) Cardiovascular effects of glucagon-like peptide 1 (GLP-1) receptor agonists. Cardiovasc Diabetol 13:142.

3. Drucker DJ (2018) Mechanisms of action and therapeutic application of glucagon-like peptide-1. Cell Metab 27(4):740–756.

4. Cho YM, Merchant CE, & Kieffer TJ (2012) Targeting the glucagon receptor family for diabetes and obesity therapy. Pharmacol Ther 135(3):247–278.

5. Samson SL & Garber A (2013) GLP-1R agonist therapy for diabetes: benefits and potential risks. Curr Opin Endocrinol Diabetes Obes 20(2):87–97.

6. Thethi TK, Pratley R, & Meier JJ (2020) Efficacy, safety and cardiovascular outcomes of once-daily oral semaglutide in patients with type 2 diabetes: The PIONEER programme. Diabetes Obes Metab 22(8):1263–1277.

7. Wright EE, Jr. & Aroda VR (2020) Clinical review of the efficacy and safety of oral semaglutide in patients with type 2 diabetes considered for injectable GLP-1 receptor agonist therapy or currently on insulin therapy. Postgrad Med:1–11.

8. Wootten D, et al. (2013) Differential activation and modulation of the glucagon-like peptide-1 receptor by small molecule ligands. Mol Pharmacol 83(4):822–834.

9. Choe HJ & Cho YM (2021) Peptidyl and non-Peptidyl oral glucagon-like peptide-1 receptor agonists. Endocrinol Metab (Seoul) 36(1):22–29.

10. Liu C, Zou Y, & Qian H (2020) GLP-1R agonists for the treatment of obesity: a patent review (2015-present). Expert Opin Ther Pat 30(10):781–794.

11. Saxena A, et al. (2020) 353-OR: Oral small molecule GLP-1R agonist PF-06882961 robustly reduces plasma glucose and body weight after 28 days in adults with T2DM. Diabetes 69(Supplement_1).

12. Zhao P, et al. (2020) Activation of the GLP-1 receptor by a non-peptidic agonist. Nature 577(7790):432–436.

13. Chen D, et al. (2007) A nonpeptidic agonist of glucagon-like peptide 1 receptors with efficacy in diabetic db/db mice. Proc Natl Acad Sci U S A 104(3):943–948.

14. Su H, et al. (2008) Boc5, a non-peptidic glucagon-like peptide-1 receptor agonist, invokes sustained glycemic control and weight loss in diabetic mice. PLoS One 3(8):e2892.

15. He M, et al. (2012) A continued saga of Boc5, the first non-peptidic glucagon-like peptide-1 receptor agonist with in vivo activities. Acta Pharmacol Sin 33(2):148–154.

16. Liu Q, et al. (2012) Cyclobutane derivatives as novel nonpeptidic small molecule agonists of glucagon-like peptide-1 receptor. J Med Chem 55(1):250–267.

17. Duan J, et al. (2020) Cryo-EM structure of an activated VIP1 receptor-G protein complex revealed by a NanoBiT tethering strategy. Nat Commun 11(1):4121.

18. Zhou F, et al. (2020) Structural basis for activation of the growth hormone-releasing hormone receptor. Nat Commun 11(1):5205.

19. Lucey M, et al. (2020) Disconnect between signalling potency and in vivo efficacy of pharmacokinetically optimised biased glucagon-like peptide-1 receptor agonists. Mol Metab 37:100991.

20. Wu F, et al. (2020) Full-length human GLP-1 receptor structure without orthosteric ligands. Nat Commun 11(1):1272.

21. Wootten D, et al. (2013) Polar transmembrane interactions drive formation of ligand-specific and signal pathway-biased family B G protein-coupled receptor conformations. Proc Natl Acad Sci U S A 110(13):5211–5216.

22. Zhang X, et al. (2020) Differential GLP-1R binding and activation by peptide and non-peptide agonists. Mol Cell 80(3):485–500.

23. Montrose-Rafizadeh C, et al. (1997) High potency antagonists of the pancreatic glucagon-like peptide-1 receptor. J Biol Chem 272(34):21201–21206.

24. Liang YL, et al. (2018) Phase-plate cryo-EM structure of a biased agonist-bound human GLP-1 receptor-Gs complex. Nature 555(7694):121–125.

25. Bueno AB, et al. (2020) Structural insights into probe-dependent positive allosterism of the GLP-1 receptor. Nat Chem Biol 16(10):1105–1110.

26. Fan H, et al. (2015) The non-peptide GLP-1 receptor agonist WB4-24 blocks inflammatory nociception by stimulating beta-endorphin release from spinal microglia. Br J Pharmacol 172(1):64–79.

27. Kawai T, et al. (2020) Structural basis for GLP-1 receptor activation by LY3502970, an orally active nonpeptide agonist. Proc Natl Acad Sci U S A 117(47):29959–29967.

28. Ma H, et al. (2020) Structural insights into the activation of GLP-1R by a small molecule agonist. Cell Res 30(12):1140–1142.

29. Zhou F, et al. (2021) Molecular basis of ligand recognition and activation of human V2 vasopressin receptor. Cell Res 31(8):929–931.

30. Cong Z, et al. (2021) Molecular insights into ago-allosteric modulation of the human glucagon-like peptide-1 receptor. Nat Commun 12(1):3763.

31. Rasmussen SG, et al. (2011) Crystal structure of the beta2 adrenergic receptor-Gs protein complex. Nature 477(7366):549–555.

32. Zheng SQ, et al. (2017) MotionCor2: anisotropic correction of beam-induced motion for improved cryo-electron microscopy. Nat Methods 14(4):331–332.

33. Zhang K (2016) Gctf: Real-time CTF determination and correction. J Struct Biol 193(1):1–12.

34. Heymann JB (2018) Guidelines for using Bsoft for high resolution reconstruction and validation of biomolecular structures from electron micrographs. Protein Sci 27(1):159–171.

35. Emsley P & Cowtan K (2004) Coot: model-building tools for molecular graphics. Acta Crystallogr D Biol Crystallogr 0(Pt 12 Pt 1):2126–2132.

36. Adams PD, et al. (2010) PHENIX: a comprehensive Python-based system for macromolecular structure solution. Acta Crystallogr D Biol Crystallogr 66(Pt 2):213–221.

37. Wu EL, et al. (2014) CHARMM-GUI Membrane Builder toward realistic biological membrane simulations. J Comput Chem 35(27):1997–2004.

38. Guvench O, et al. (2011) CHARMM additive all-atom force field for carbohydrate derivatives and its utility in polysaccharide and carbohydrate-protein modeling. J Chem Theory Comput 7(10):3162–3180.

39. Vanommeslaeghe K, et al. (2010) CHARMM general force field: A force field for drug-like molecules compatible with the CHARMM all-atom additive biological force fields. J Comput Chem 31(4):671–690.

40. Hess B (2008) P-LINCS: A parallel linear constraint solver for molecular simulation. J Chem Theory Comput 4(1):116–122.

41. Aoki KM & Yonezawa F (1992) Constant-pressure molecular-dynamics simulations of the crystal-smectic transition in systems of soft parallel spherocylinders. Phys Rev A 46(10):6541–6549.

