## Supporting Information for "Structural basis of peptidomimetic agonism revealed by small molecule GLP-1R agonists Boc5 and WB4-24"

^a^Department of Pharmacology, School of Basic Medical Sciences, Fudan University, Shanghai 200032, China; ^b^Department of Biophysics and Department of Pathology of Sir Run Run Shaw Hospital, Zhejiang University School of Medicine, Hangzhou 310058, China; ^c^School of Pharmacy, Fudan University, Shanghai 201203, China; ^d^School of Artificial Intelligence and Automation, Huazhong University of Science and Technology, Wuhan 430074, China; ^e^The National Center for Drug Screening, Shanghai Institute of Materia Medica, Chinese Academy of Sciences, Shanghai 201203, China; ^f^University of Chinese Academy of Sciences, Beijing 100049, China; ^g^The CAS Key Laboratory of Receptor Research, Shanghai Institute of Materia Medica, Chinese Academy of Sciences, Shanghai 201203, China; ^h^Research Center for Deepsea Bioresources, Sanya, Hainan 572025, China.

^1^Z.C. and Q.Z. contributed equally to this work.

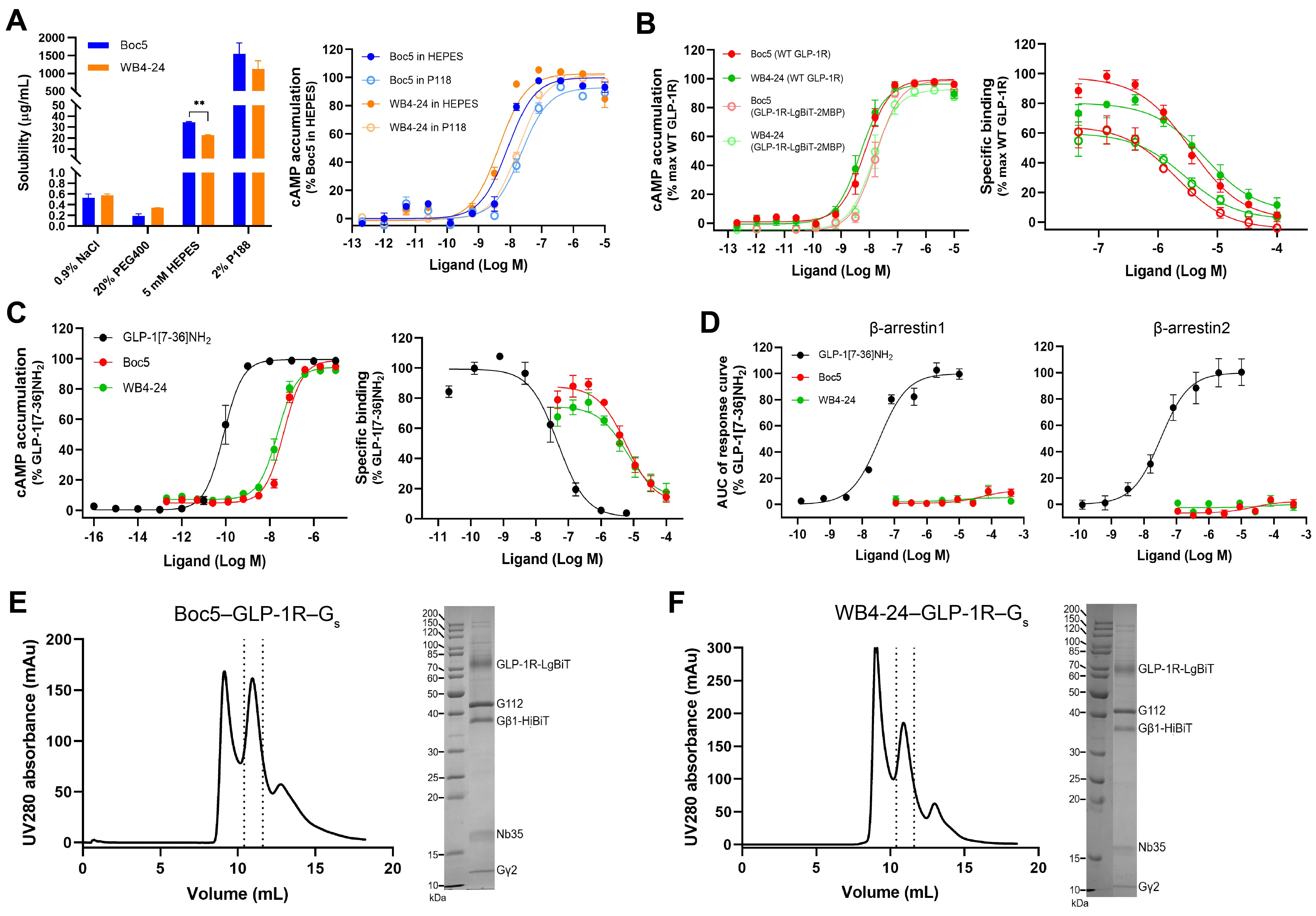

**Fig. S1. Characterization of non-peptidic GLP-1R agonists and purification of Boc5- and WB4-24-bound GLP-1R–G_s_ complexes.** (*A*) Solubility of Boc5 and WB4-24 in different solvents (left). The surfactant P188 had no significant influence on Boc5 and WB4-24 induced receptor activation (right). (*B*) cAMP responses following Boc5 and WB4-24 stimulation in HEK293T cells transfected with the wild-type (WT) or modified GLP-1R constructs (left). Binding of Boc5 and WB4-24 to the WT or modified GLP-1R in competition with ^125^I-GLP-1 (right). (*C*, *D*) GLP-1, Boc5 and WB4-24 induced cAMP accumulation (*C*) and β-arrestin 1/2 recruitment (*D*). (*E*, *F*) Size-exclusion chromatography elution profile and corresponding SDS-PAGE gel of the Boc5–GLP-1R–G_s_–Nb35 (*E*) and WB4-24–GLP-1R–G_s_–Nb35 (*E*) complexes. G112 is an engineered Gα_s_ protein. Data are shown as means ± S.E.M. from at least three independent experiments.

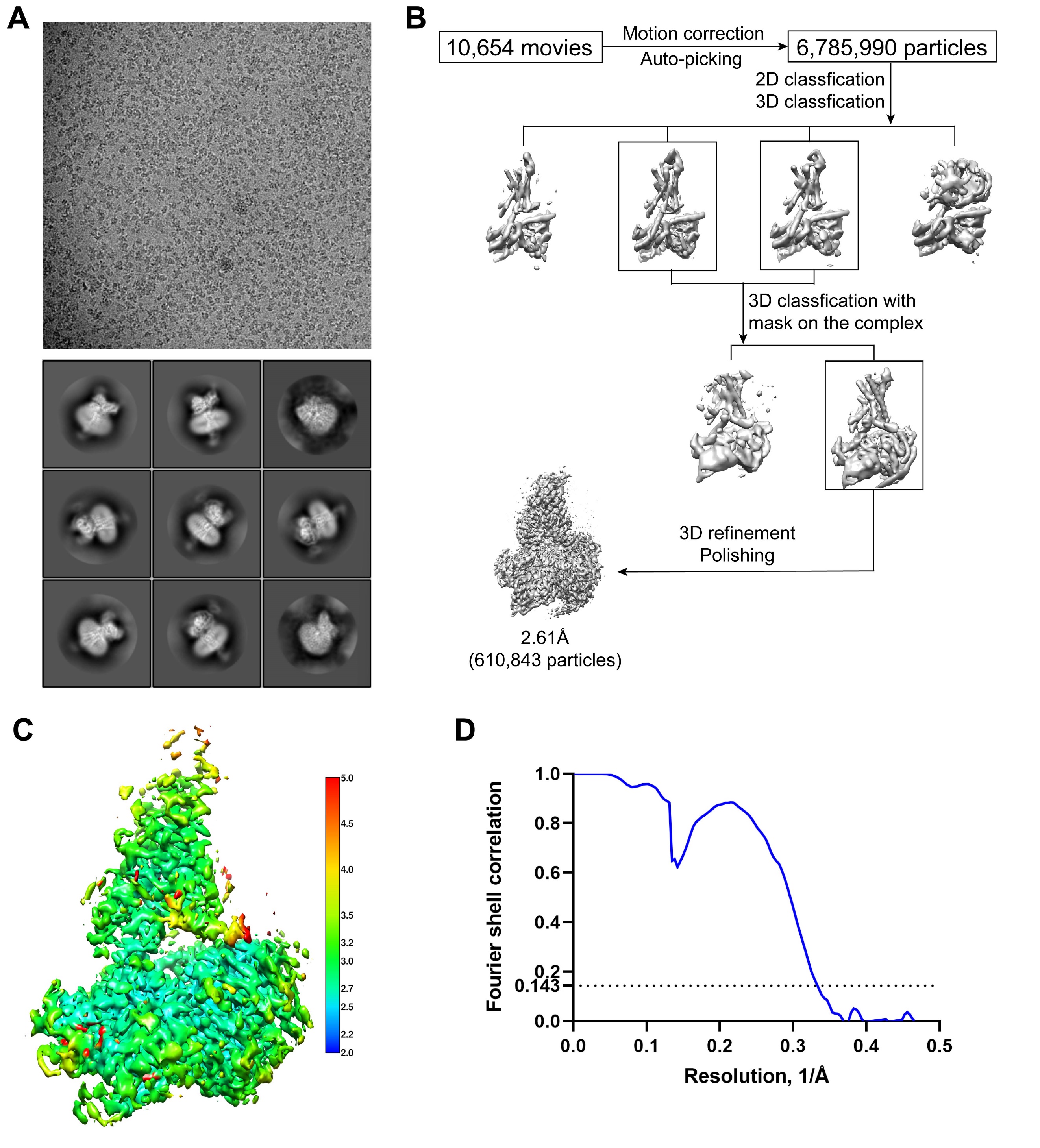
**Fig. S2. Cryo-EM data processing and validation of the Boc5–GLP-1R–G_s_–Nb35 complex.** (*A*) Representative cryo-EM micrograph and two-dimensional class averages. (*B*) Flow chart of cryo-EM data processing. (*C*) Local resolution distribution map of the complex. (*D*) Fourier shell correlation (FSC) curve of the overall refined receptor.

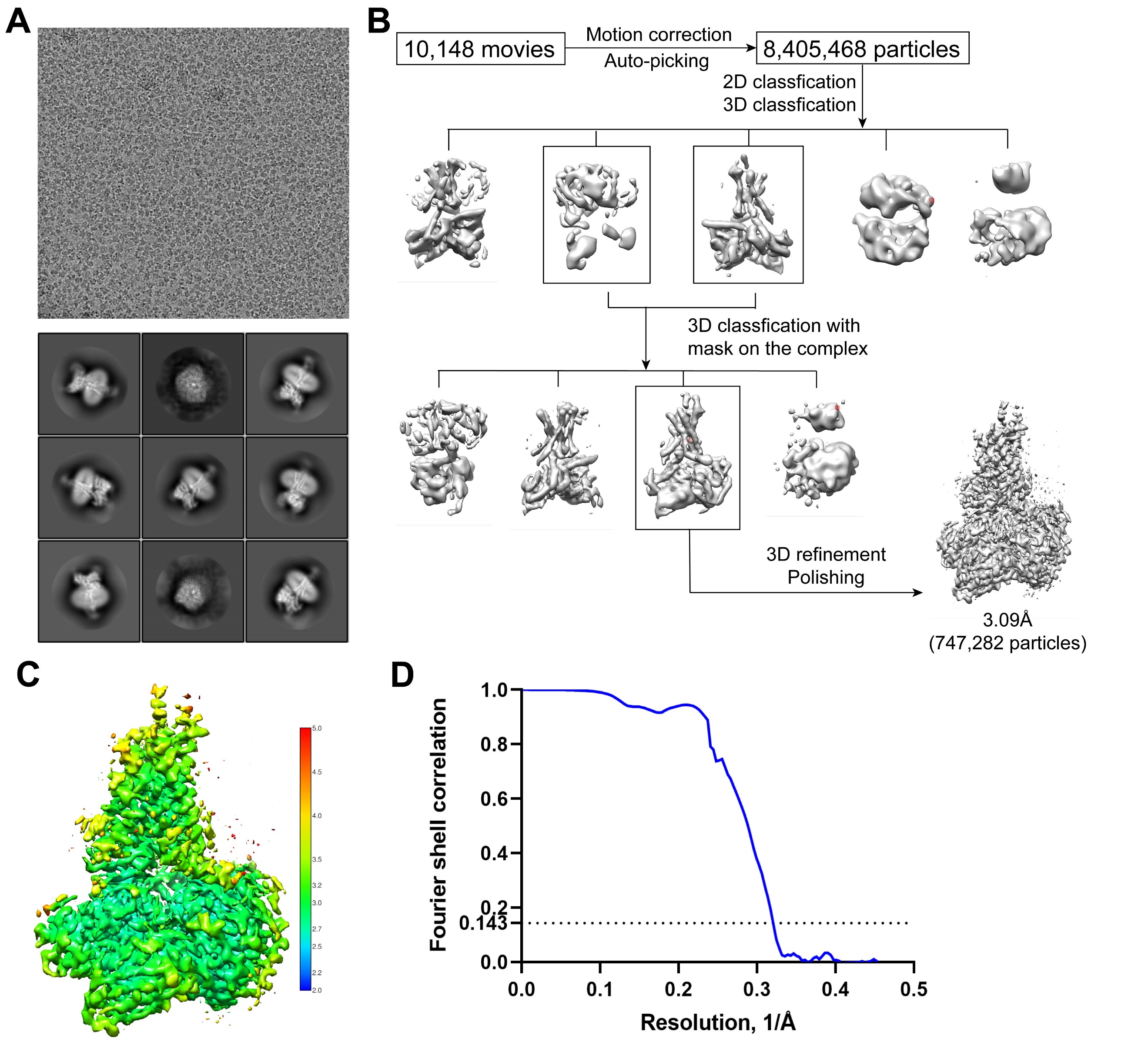

**Fig. S3. Cryo-EM data processing and validation of the WB4-24–GLP-1R–G_s_–Nb35 complex.** (*A*) Representative cryo-EM micrograph and two-dimensional class averages. (*B*) Flow chart of cryo-EM data processing. (*C*) Local resolution distribution map of the complex. (*D*) FSC curves of the overall refined receptor.

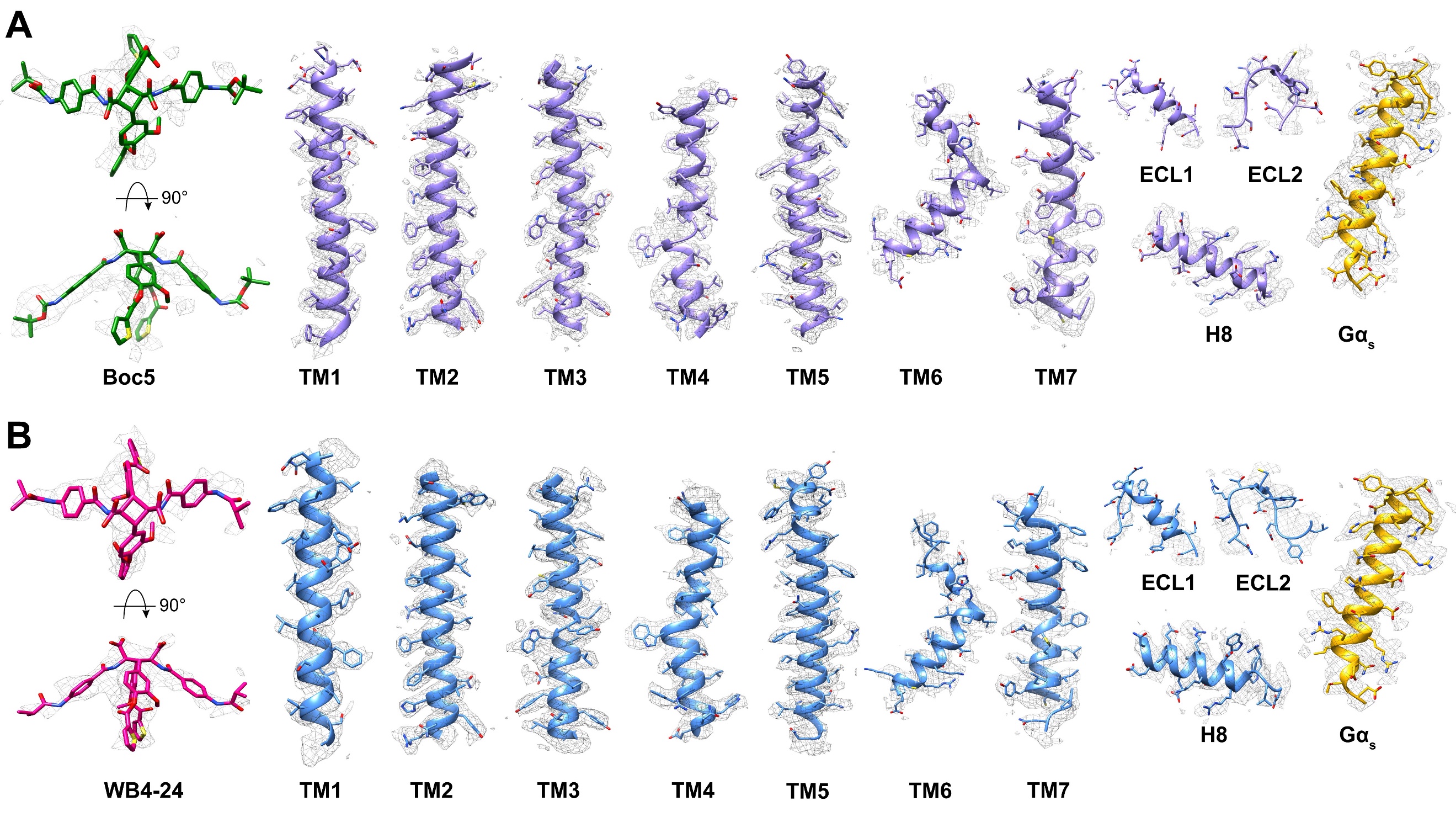

**Fig. S4. Cryo-EM density maps of the Boc5- and WB4-24-bound GLP-1R–G_s_ structures.** (*A*) Cryo-EM density map and model of the Boc5–GLP-1R–G_s_ structure are shown for Boc5, all seven-transmembrane (TM) α-helices, ECL1, ECL2, helix 8 (H8) of GLP-1R, and α5-helix of Gα_s_. (*B*) Cryo-EM density map and model of the WB4-24–GLP-1R–G_s_ structure are shown for WB4-24, all TM α-helices, ECL1, ECL2, H8 of GLP-1R, and α5-helix of Gα_s_.

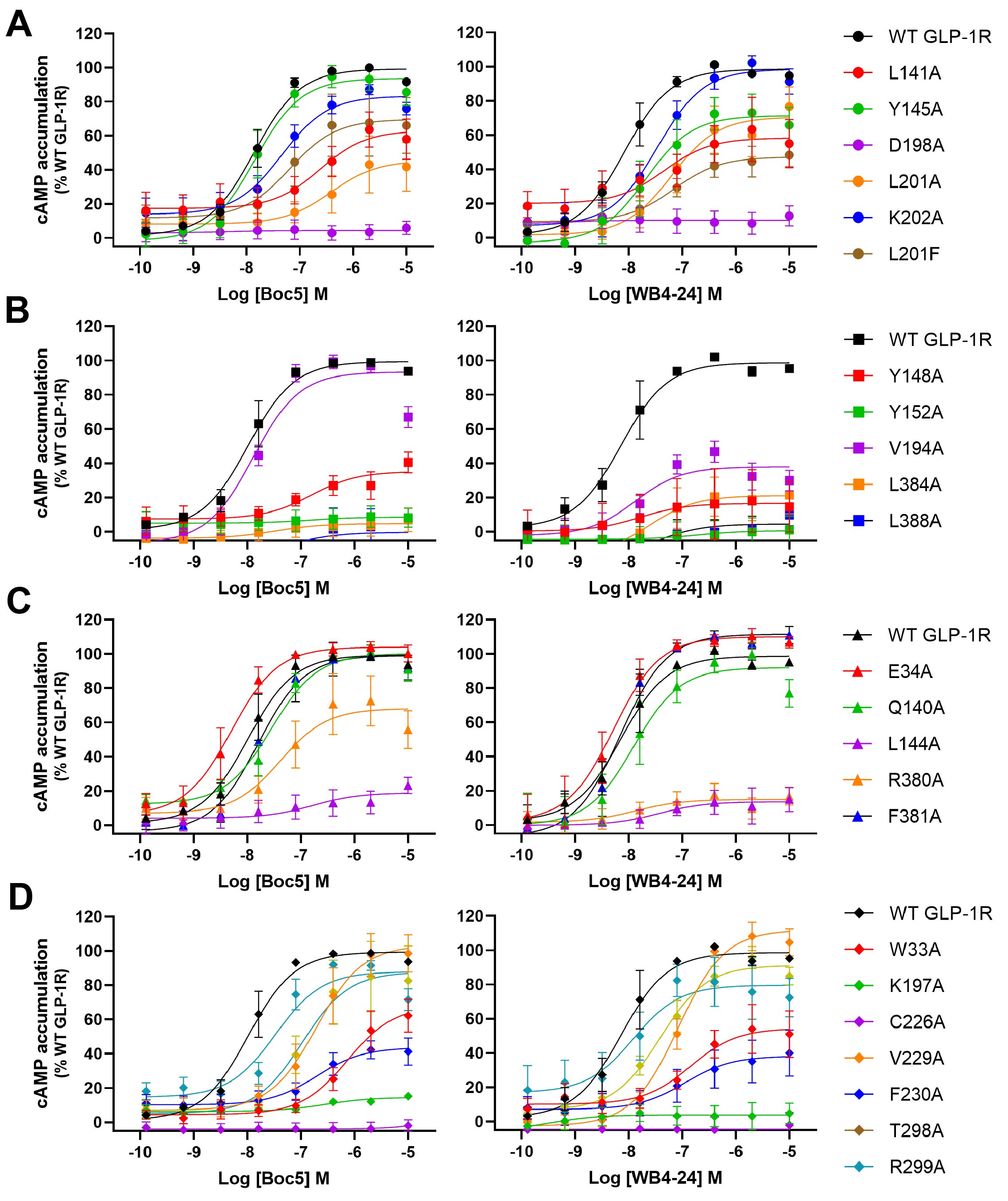

**Fig. S5. Ligand mediated cAMP accumulation at wild-type (WT) or mutant GLP-1R.** Concentration-response curves for alanine mutation of residues interacting with arms A1-1 (*A*), A1-2 (*B*), B1-1 or B2-1 (*C*), and B2-1 or B2-2 (*D*) of Boc5 and WB4-24. Data are shown as means ± S.E.M. of at least four independent experiments performed in quadruplicate.

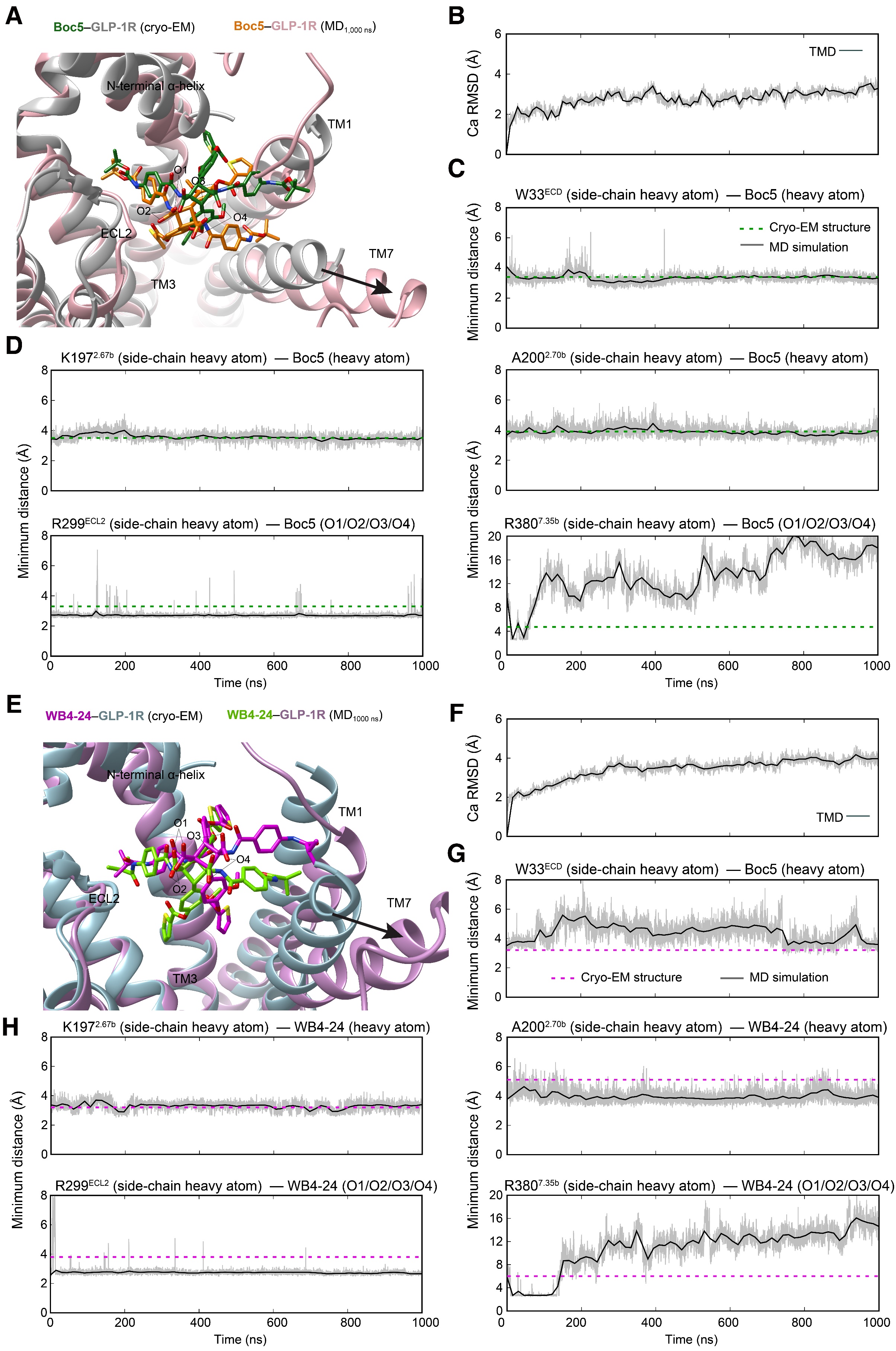

**Fig. S6. Molecular dynamics (MD) simulations of Boc5 or WB4-24-bound active GLP-1R.** (*A*) Comparison of the Boc5 conformation between simulation snapshot and the cryo-EM structure. (*B*) Root mean square deviation (RMSD) of Cα positions of the GLP-1R TMD, where all snapshots were superimposed on the cryo-EM structure of Boc5-bound GLP-1R TMD using the Cα atoms. (*C*) Minimum distance between the heavy atoms of Boc5 and the ECD residue W33 suggests that Boc5 steadily interacts with the N-terminal α-helix of ECD. (*D*) Analysis of the MD simulation trajectories in (*A*): representative minimum distances between the heavy atoms of Boc5 and TMD residues (top left, K197^2.67b^－Boc5; top right, A200^2.70b^－Boc5; bottom left, R299^ECL2^－O1/O2/O3/O4 in Boc5; bottom right, R380^7.45b^－O1/O2/O3/O4 in Boc5). (*E*) Comparison of the WB4-24 conformation between simulation snapshot and the cryo-EM structure. (*F*) RMSD of Cα positions of the GLP-1R TMD, where all snapshots were superimposed on the cryo-EM structure of WB4-24-bound GLP-1R TMD using the Cα atoms. (*G*) Minimum distance between the heavy atoms of WB4-24 and the ECD residue W33. (*H*) Analysis of the MD simulation trajectories in (*E*): representative minimum distances between the heavy atoms of WB4-24 and TMD residues (top left, K197^2.67b^－WB4-24; top right, A200^2.70b^－WB4-24; bottom left, R299^ECL2^－O1/O2/O3/O4 in WB4-24; bottom right, R380^7.45b^－O1/O2/O3/O4 in WB4-24). The thick and thin traces represent moving averages and original, unsmoothed values, respectively.

**Table S1. Cryo-EM data collection, model refinement and validation statistics.**

| **Data collection and processing** | **Boc5-GLP-1R-G_s_-Nb35** | **WB4-24-GLP-1R-G_s_-Nb35** |
| --- | --- | --- |
| Magnification | 130,000 | 130,000 |
| Voltage (kV) | 300 | 300 |
| Electron exposure (e–/Å^2^) | 80 | 80 |
| Defocus range (μm) | -1.2 to -2.2 | -1.2 to -2.2 |
| Pixel size (Å) | 1.071 | 1.071 |
| Symmetry imposed | C1 | C1 |
| Initial particle image (no.) | 6,785,990 | 8,405,468 |
| Final particle image (no.) | 610,843 | 747,282 |
| Map resolution (Å)  FSC threshold | 2.61  0.143 | 2.60  0.143 |
| Map resolution range (Å) | 2.5-4.9 | 2.2-4.0 |
| **Refinement** |  |  |
| Initial model used (PDB code) | 6X19 | 6X19 |
| Model resolution (Å)  FSC threshold | 2.7  0.5 | 3.03  0.5 |
| Map sharpening B factor (Å^2^) | -82.56 | -82.56 |
| Model composition  Non-hydrogen atom  Protein residue | 8493  1051 | 8047  1019 |
| B factors (Å^2^)  Protein | 46.54 | 93.64 |
| Root mean square deviation  Bond length (Å)  Bond angle (°) | 0.003  0.642 | 0.004  0.679 |
| Validation  MolProbity score  Clash score  Poor rotamer (%) | 1.59  7.32  0.11 | 1.39  3.50  0.0 |
| Ramachandran plot  Favored (%)  Allowed (%)  Disallowed (%) | 96.90  3.10  0.0 | 96.31  3.69  0.0 |

**Table S2.** ***In vitro* pharmacology of GLP-1, Boc5 and WB4-24 at wild-type and mutant GLP-1R constructs.**

|  | **cAMP** | | **Binding** | **β-arrestin 1** | | **β-arrestin 2** | |
| --- | --- | --- | --- | --- | --- | --- | --- |
|  | pEC_50_ | Emax | pKi | pEC_50_ | Emax | pEC_50_ | Emax |
| **WT GLP-1R** | | | | | | | |
| GLP-1 | 10.09±0.06 | 99.53±1.67 | 7.44±0.14 | 7.47±0.09 | 100±3.01 | 7.48±0.14 | 100±4.71 |
| Boc5 | 7.35±0.05 | 99.06±1.86 | 5.50±0.11 | 4.30±0.41 | 10.91±2.78 | 4.64±0.49 | 2.76±2.54 |
| WB4-24 | 7.64±0.05 | 94.47±1.67 | 5.40±0.12 | 5.02±1.56 | 5.26±2.56 | NA | NA |
| **GLP-1R-LgBiT-2MBP** | | | | | | | |
| GLP-1 | ND | ND | ND | ND | ND | ND | ND |
| Boc5 | 7.78±0.07 | 98.94±2.63 | 5.99±0.11 | ND | ND | ND | ND |
| WB4-24 | 7.87±.08 | 92.59±2.75 | 6.00±0.09 | ND | ND | ND | ND |

The functional potency (EC_50_) and (E_max_) values for cAMP accumulation and β-arrestin recruitment assays were analyzed using a three-parameter logistic equation. pEC_50_ is the negative logarithm of the molar concentration of agonist that induced half the maximal response. E_max_ is expressed as a percentage of the response induced by GLP-1. Binding data were analyzed using a three-parameter logistic equation and normalized to the maximal binding of ^125^I-GLP-1. pKi is the negative logarithm of peptide affinity. All values are means ± S.E.M. of at least three independent experiments conducted in quadruplicate (cAMP accumulation) or duplicate (binding assay and β-arrestin recruitment). WT, wild-type; ND, not determined; NA, not active.

**Table S3. Effects of residue mutation in the ligand binding pocket on cAMP accumulation.**

| **GLP-1R residue** | **Boc5** | | **WB4-24** | |
| --- | --- | --- | --- | --- |
|  | pEC_50_ | E_max_ | pEC_50_ | E_max_ |
| **Interaction with A-1** | | | | |
| WT GLP-1R | 7.86±0.09 | 99.28±2.63 | 8.08±0.10 | 98.57±2.92 |
| L141^1.36b^A | 6.63±0.39** | 63.12±8.05** | 7.34±0.54** | 58.35±7.23** |
| Y145^1.40b^A | 7.84±0.16 | 93.75±4.12 | 7.68±0.29* | 71.41±7.01** |
| D198^2.68b^A | NA** | 4.43±2.33** | NA** | 10.28±2.97** |
| L201^2.71b^A | 6.42±0.49** | 45.48±8.98** | 7.20±0.25** | 70.63±6.68** |
| K202^2.72b^A | 7.34±0.17* | 83.63±4.25* | 7.47±0.15* | 98.52±4.88 |
| **Interaction with A-2** | | | | |
| Y148^1.43b^A | 6.81±0.41** | 35.29±4.82** | NA** | 16.66±7.94** |
| Y152^1.47b^A | 6.82±2.86** | 8.66±4.22** | NA** | 0.69±6.41** |
| V194^2.64b^A | 7.98±0.15 | 93.34±4.31 | 7.88±0.31 | 38.04±3.82** |
| L384^7.39b^A | 7.47±1.06* | 4.85±3.17** | 7.67±0.45* | 21.24±5.30** |
| L388^7.43b^A | 7.12±1.76 | 0.28±8.16** | 7.49±0.71 | 4.64±4.93** |
| **Interaction with B1-1 or B2-1** | | | | |
| E34^ECD^A | 8.30±0.10* | 104.02±3.78 | 8.28±0.16 | 110.06±4.67 |
| Q140^1.35b^A | 7.55±0.12 | 100.29±3.42 | 7.91±0.24 | 92.05±6.35 |
| L144^1.39b^A | 6.78±0.81** | 18.97±5.12** | 7.44±0.73 | 13.68±3.50** |
| R380^7.35b^A | 7.38±0.33* | 68.18±7.12** | 7.87±0.76 | 15.01±3.12** |
| F381^7.36b^A | 7.80±0.14 | 99.14±4.66 | 8.16±0.15 | 111.53±5.04 |
| **Interaction with B1-2 or B2-2** | | | | |
| W33^ECD^A | 6.13±0.21** | 67.98±7.52** | 6.39±0.40** | 54.48±6.48** |
| K197^2.67b^A | 6.55±0.47** | 14.59±1.82** | NA** | 3.95±2.42** |
| C226^3.29b^A | NA** | 3.73±2.46** | NA** | NA** |
| V229^3.32b^A | 6.71±0.19** | 103.60±8.02 | 7.34±0.18** | 111.90±4.27 |
| F230^3.33b^A | 6.71±0.32** | 43.74±4.70** | 6.96±0.58** | 38.29±7.02** |
| T298^ECL2^A | 7.48±0.18* | 87.94±4.55* | 7.92±0.38 | 79.69±7.24** |
| R299^ECL2^A | 6.98±0.27** | 87.81±9.09 | 7.54±0.18* | 91.23±3.49 |

cAMP accumulation data were analyzed using a three-parameter logistic equation to determine pEC_50_ and E_max_ values. pEC_50_ is the negative logarithm of the molar concentration of agonist that induced half the maximal response. E_max_ for mutants is expressed as a percentage of the response induced by the wild-type (WT) GLP-1R. All values are means ± S.E.M. of at least three independent experiments conducted in quadruplicate. One-way ANOVA was used to determine statistical significance (**P<0.01, *P<0.05). NA, not active.

**Table S4. Interaction between GLP-1R and Boc5 or WB4-24.**

| **GLP-1R residue** | **Boc5** | | **WB4-24** |
| --- | --- | --- | --- |
| W33^ECD^ | | Stacking interaction | Stacking interaction |
| Q140^1.35b^ | | Hydrophobic contacts | Hydrophobic contacts |
| L141^1.36b^ | | Hydrophobic contacts | Hydrophobic contacts |
| L144^1.39b^ | | Hydrophobic contacts | Hydrophobic contacts |
| Y145^1.40b^ | | Stacking interaction | Stacking interaction |
| Y148^1.43b^ | | Hydrophobic contacts | Hydrophobic contacts |
| Y152^1.47b^ | | Hydrophobic contacts | Hydrophobic contacts |
| R190^2.60b^ | | Stacking interaction | Stacking interaction |
| V194^2.64b^ | | Hydrophobic contacts | Hydrophobic contacts |
| I196^2.66b^ | | Hydrophobic contacts |  |
| K197^2.67b^ | | Stacking interaction  Hydrophobic contacts | Stacking interaction  Hydrophobic contacts |
| D198^2.68b^ | |  | Hydrophobic contacts |
| A200^2.70b^ | | Hydrophobic contacts |  |
| L201^2.71b^ | | Hydrophobic contacts | Hydrophobic contacts |
| K202^2.72b^ | | Hydrophobic contacts | Hydrophobic contacts |
| C226^3.29b^ | | Hydrophobic contacts | Hydrophobic contacts |
| V229^3.32b^ | | Hydrophobic contacts | Hydrophobic contacts |
| F230^3.33b^ | | Stacking interaction | Stacking interaction |
| M233^3.36b^ | | Hydrophobic contacts |  |
| T298^ECL2^ | | Hydrogen bond | Hydrogen bond |
| R299^ECL2^ | | Salt bridge | Salt bridge |
| R380^7.35b^ | | Salt bridge | Salt bridge |
| F381^7.36b^ | | Hydrophobic contacts | Hydrophobic contacts |
| L384^7.39b^ | | Hydrophobic contacts | Hydrophobic contacts |
| F385^7.40b^ | |  | Hydrophobic contacts |
| L388^7.43b^ | | Hydrophobic contacts | Hydrophobic contacts |

**Table S5. Comparison of *in vivo* and *in vitro* activities of peptidic and non-peptidic GLP-1R agonists.**

| **Ligand** | **Peptide** | **Small molecule** | | | | |
| --- | --- | --- | --- | --- | --- | --- |
|  | **GLP-1** | **Boc5** | **WB4-24** | **TT-OAD2** | **LY3502970** | **PF-06882961** |
| ***In vitro* activity** | | | | | | |
| cAMP response | Full agonist | Full agonist | Full agonist | Partial agonist | Partial agonist | Full agonist |
| β-arrestin recruitment | Full agonist | NA | NA | NA | NA | Partial agonist |
| ***In vivo* activity** | | | | | | |
| Insulin release | + | + | + | + | + | + |
| Appetite reduction | + | + | + | + | + | + |
| HbA1c reduction | + | + | + | + | + | + |
| Body weight reduction | + | + | + | ? | ? | + |
| CVD benefit | + | + | + | ? | ? | + |

HbA1c, hemoglobin A1c; CVD, cardiovascular disease.
